# Loss of CHCHD2 and CHCHD10 reveals differential vulnerability to bioenergetic failure in cardiac and skeletal muscle

**DOI:** 10.64898/2026.07.15.738671

**Authors:** Jule Gerlach, Juan Diego Hernández-Camacho, Hendrik Nolte, Frank Chenfei Ning, Joana Filipa Silva-Rodrigues, David Alsina, Dusanka Milenkovic, Jelena Misic, Annelie Falkevall, Thomas Langer, Timothy Wai, Roberta Filograna

## Abstract

Mutations in the mitochondrial proteins CHCHD2 and CHCHD10 cause severe neurodegenerative and neuromuscular disorders, yet their physiological functions remain poorly defined. CHCHD2 and CHCHD10 localize to the mitochondrial intermembrane space, where they assemble into a high-molecular-weight complex. Here, we generated *Chchd2/Chchd10* double-knockout (DKO) mice to define the *in vivo* role of this complex. DKO mice developed reduced lean mass, progressive muscle weakness, and oxidative phosphorylation defects in both cardiac and skeletal muscle. Despite comparable bioenergetic impairment, CHCHD2-CHCHD10 deficiency elicited tissue-specific responses distinct from those induced by the disease-associated *CHCHD10 S59L* mutation. The heart accumulated enlarged mitochondria with disrupted ultrastructure and underwent adaptive proteomic remodeling that preserved myocardial contractility into adulthood. By contrast, skeletal muscle exhibited limited proteomic alterations, accompanied by profound changes in lipid composition, reduced expression of the myogenic regulators, and altered myofiber size. Mechanistically, depletion of CHCHD2 and CHCHD10 in primary satellite cells impaired proliferation and myogenic differentiation, suggesting that defective postnatal myogenesis contributes to the muscle growth defect in DKO mice.

Together, these findings identify the CHCHD2-CHCHD10 complex as a critical regulator of mitochondrial integrity and demonstrate that its loss drives tissue-specific defects culminating in diverse pathological outcomes.

## Introduction

Mitochondria sustain cellular physiology by coordinating essential processes, including energy production, metabolic regulation, cell death signaling, and stress responses^1^. Central to these functions is the mitochondrial intermembrane space (IMS), a highly specialized compartment positioned between the outer mitochondrial membrane (OMM), which interfaces with the cytosol, and the inner mitochondrial membrane (IMM), whose highly folded cristae house the bulk of the oxidative phosphorylation (OXPHOS) machinery^2,3^. This strategic location enables the IMS to mediate communication between mitochondria and the cytosol^2^, integrating the regulation of mitochondrial protein import, proteostasis and signaling pathways.

The IMS hosts multiple quality control and surveillance mechanisms required for the maintenance and remodeling of the organelle during metabolic alterations^3^. These include IMM-bound proteases with IMS-facing catalytic domains that regulate protein turnover, mitochondrial dynamics, and stress responses. A key example is the i-AAA protease YME1L, which plays a central role in mitochondrial proteostasis by degrading misfolded proteins and components of the import machinery^4,5^. It also regulates phospholipid metabolism^4^ and mitochondrial dynamics through the processing of substrates such as Optic Atrophy 1 (OPA1), a dynamin-like GTPase that performs IMM fusion^3,6^. Consistent with its essential housekeeping functions, loss of YME1L results in embryonic lethality in mice^6^.

By contrast, the protease OMA1 is activated in response to mitochondrial stress and functions as both a proteostasis regulator and a signaling node^7–9^. Upon activation, OMA1 cleaves OPA1^10^, promoting mitochondrial fragmentation. In parallel, it processes DELE1, thereby triggering the integrated stress response (ISR) and driving broader cellular adaptations, including attenuation of cytosolic translation and metabolic rewiring^11,12^. *In vivo* studies reveal a complex picture in which OMA1-dependent pathways regulate cell survival in a tissue- and stress-specific manner^13^, with OMA1 ablation having either beneficial^14^ or detrimental^15,16^ effects depending on the context. These examples illustrate the functional diversity of IMS pathways, encompassing both constitutive and stress-responsive programs that preserve mitochondrial integrity and adapt to changing cellular demands. Despite these advances, the physiological functions of many other IMS proteins remain poorly understood. Among these are the paralogous proteins CHCHD2 and CHCHD10, which have emerged as important contributors to human disease^17^.

Mutations in *CHCHD2* are primarily associated with Parkinson’s disease^18,19^, whereas variants in *CHCHD10* are linked to a broader spectrum of disorders, including amyotrophic lateral sclerosis (ALS), frontotemporal dementia (FTD)^20^, spinal muscular atrophy^21^, cardiomyopathy, and myopathy^22^. Several disease-associated variants are thought to act through toxic gain-of-function mechanisms, involving the accumulation of aberrant protein species that ultimately impair mitochondrial function^23^. In humans, the well-characterised *CHCHD10 p.S59L* mutation causes a multisystemic disorder marked by mitochondrial dysfunction and pronounced tissue-specific vulnerability. Key features of this phenotype are recapitulated in heterozygous *Chchd10 S55L* knock-in mice (equivalent to the S59L human mutation), whereas they are absent in *Chchd10* knockout models, supporting a toxic gain-of-function mechanism^24,25^. In affected tissues, OMA1 activation, aberrant OPA1 processing, and a robust ISR have been observed and contribute to disease pathology^26^.

CHCHD2 and CHCHD10 are small (∼15 kDa) CHCH-domain proteins that share high sequence identity and physically interact to form a stable ∼150-200 kDa protein complex^27,28^. Although they have been associated with mitochondrial cristae organization^29^, respiratory chain function^30^, and mitochondrial dynamics^31^, the physiological function of the CHCHD2-CHCHD10 complex remains unresolved. We previously showed that whole-body *Chchd2* knockout mice develop mild motor deficits and reduced striatal dopamine levels, despite preserved OXPHOS capacity and no overt loss of dopaminergic neurons^32^. Notably, CHCHD10 retains its ability to assemble into high-molecular-weight oligomers in the absence of CHCHD2, indicating partial functional compensation. The abundance and assembly of the CHCHD2-CHCHD10 complex are dynamically regulated by mitochondrial fitness across different tissues^32^, and *in vitro* CHCHD2 expression confers protection against mitochondrial damage^32,33^. These observations raise the hypothesis that the CHCHD2-CHCHD10 complex operates as a stress-response factor critical for supporting cell survival^34^.

To directly define the *in vivo* functions of CHCHD2 and CHCHD10 and address potential redundancy, we generated whole-body *Chchd2/Chch10* double knockout (DKO) mice. We show that loss of this protein complex results in substantial bioenergetic failure and leads to marked, tissue-specific defects in cardiac and skeletal muscle. Our results show that the CHCHD2-CHCHD10 complex is required to maintain mitochondrial homeostasis in adult muscle tissues and point to its involvement in previously unrecognized IMS-based mechanisms coordinating tissue-specific responses to mitochondrial stress.

## Results

### Loss of CHCHD2 and CHCHD10 impairs lean-mass growth and leads to muscle weakness

To explore the role of the CHCHD2-CHCHD10 complex in mammalian mitochondria, we generated *Chchd2/Chchd10* DKO mice by crossing previously described *Chchd2* KO (D2 KO) mice^32^ with newly generated CRISPR-Cas9-mediated *Chchd10* KO (D10 KO) mice. Western blot analyses confirmed the absence of CHCHD2 and CHCHD10 in DKO mouse tissues (Supplementary Fig. 1a). Whole-body homozygous DKO mice were viable but modestly underrepresented relative to expected Mendelian ratios, with no detectable sex bias (Supplementary Fig. 1b).

To assess the impact of CHCHD2 and CHCHD10 ablation on the gross phenotype, body weight was monitored from 3 to 12 months of age. Male DKO mice were smaller and exhibited reduced weight gain from 3 months of age, whereas female DKO mice only displayed a significant reduction at 12 months (Fig.1a). To determine whether these differences reflected altered body composition, lean and fat mass were quantified by EchoMRI from 3 months of age. Both male and female DKO mice showed decreased lean mass at 6 and 12 months compared with control and D10 KO mice, whereas fat mass remained largely unchanged (Fig.1 b).

**Fig. 1:**
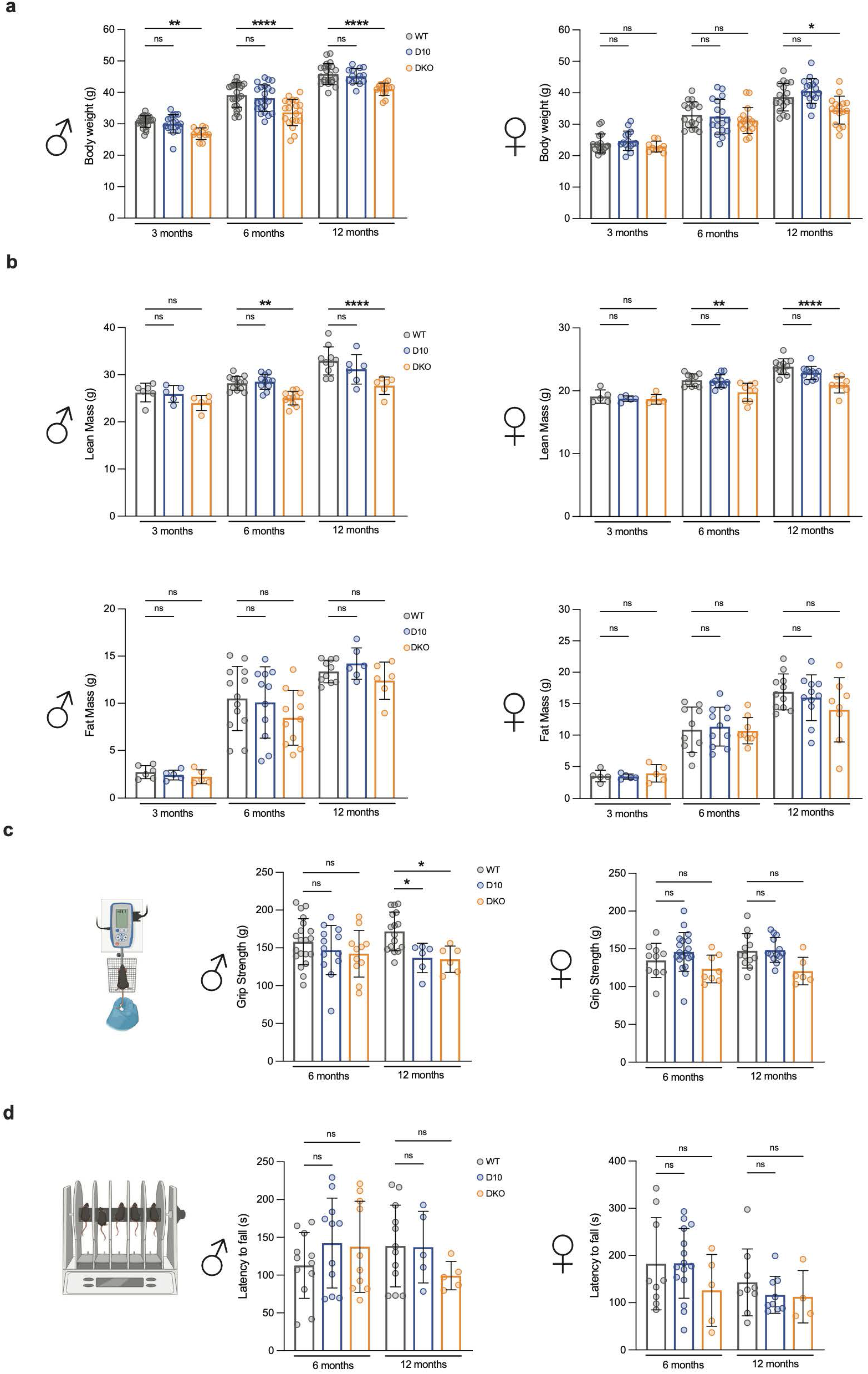
DKO mice exhibit reduced body weight, decreased lean mass, and progressive decline in muscle strength. **a,** Body weight (BW) of male and female Wildtype (WT), *Chchd10^-/-^* (D10 KO), and *Chchd2^-/-^ and Chchd10^-/-^* (DKO) mice at 3, 6, and 12 months (n > 14 per genotype and time point). **b,** Lean and fat mass, measured by echoMRI in male and female WT, D10 KO, and DKO mice at 3, 6, and 12 months (n> 5 per genotype and time point). **c,** Muscle strength (g) measured by grip strength, in male and female mice at 6 and 12 months. **d,** Motor coordination and endurance assessed by latency to fall (s) from the Rotarod in male and female mice at 6 and 12 months (n> 4 per genotype and time point). Data in **a-d** are presented as means ± SD. *p < 0.05; **p < 0.01; ****p < 0.0001; ns, not significant.

To investigate the functional consequences of these skeletal muscle deficits, motor performance tests were conducted to assess muscle strength and coordination. Four-limb grip measurements revealed reduced absolute grip strength in male D10 KO and DKO mice at 12 months of age. Female DKO mice manifested a similar but non-significant trend at 6 and 12 months (Fig.1c). Rotarod analyses indicated a modest but not statistically significant decline in coordination and endurance in both male and female DKO mice (Fig. 1d; Supplementary Fig. 1c).

Together, these data show that DKO mice develop a muscle phenotype characterized by reduced lean mass and mild muscle weakness, with more pronounced effects observed in males.

### CHCHD2 and CHCHD10 ablation impairs OXPHOS capacity in cardiac and skeletal muscle

Next, we examined how the loss of the CHCHD2-CHCHD10 complex affects OXPHOS capacity in cardiac and skeletal muscle. Spectrophotometric measurements of respiratory chain complexes in mitochondria isolated from 6-month-old hearts revealed reduced activities of complexes II, I+III, and IV in DKO mice, with the most severe decrease (∼40%) observed in complex I. These deficiencies were slightly exacerbated at 12 months, suggesting a slow, progressive decline in cardiac OXPHOS capacity (Fig.2a).

In skeletal muscle, specifically in mitochondria isolated from quadriceps femoris, OXPHOS defects were more profound. Marked reductions in complexes I (∼60%), I+III, and IV (∼30%) activities were already present at 6 months. These alterations showed a modest progression with age, with complex IV showing the greatest reduction (∼45%) at 12 months (Fig. 2a). To determine whether the observed functional impairments were associated with defective complex assembly or maintenance, we performed in-gel activity assays for complexes I and IV. Under digitonin solubilization conditions, both complexes were only modestly affected in DKO heart and quadriceps, with no evidence of defective assembly of individual complexes (Fig. 2b). This was further supported by analyses of mitochondria from 12-month-old quadriceps solubilized using both mild (digitonin) and harsh (n-dodecyl-β-D-maltoside, DDM) detergents, which showed preserved activity of individual complexes but a mild reduction in supercomplex-associated activity (Supplementary Fig. 2a). This pattern suggests that DKO mice may exhibit alterations in the IMM that destabilize higher-order respiratory chain assemblies.

**Fig. 2:**
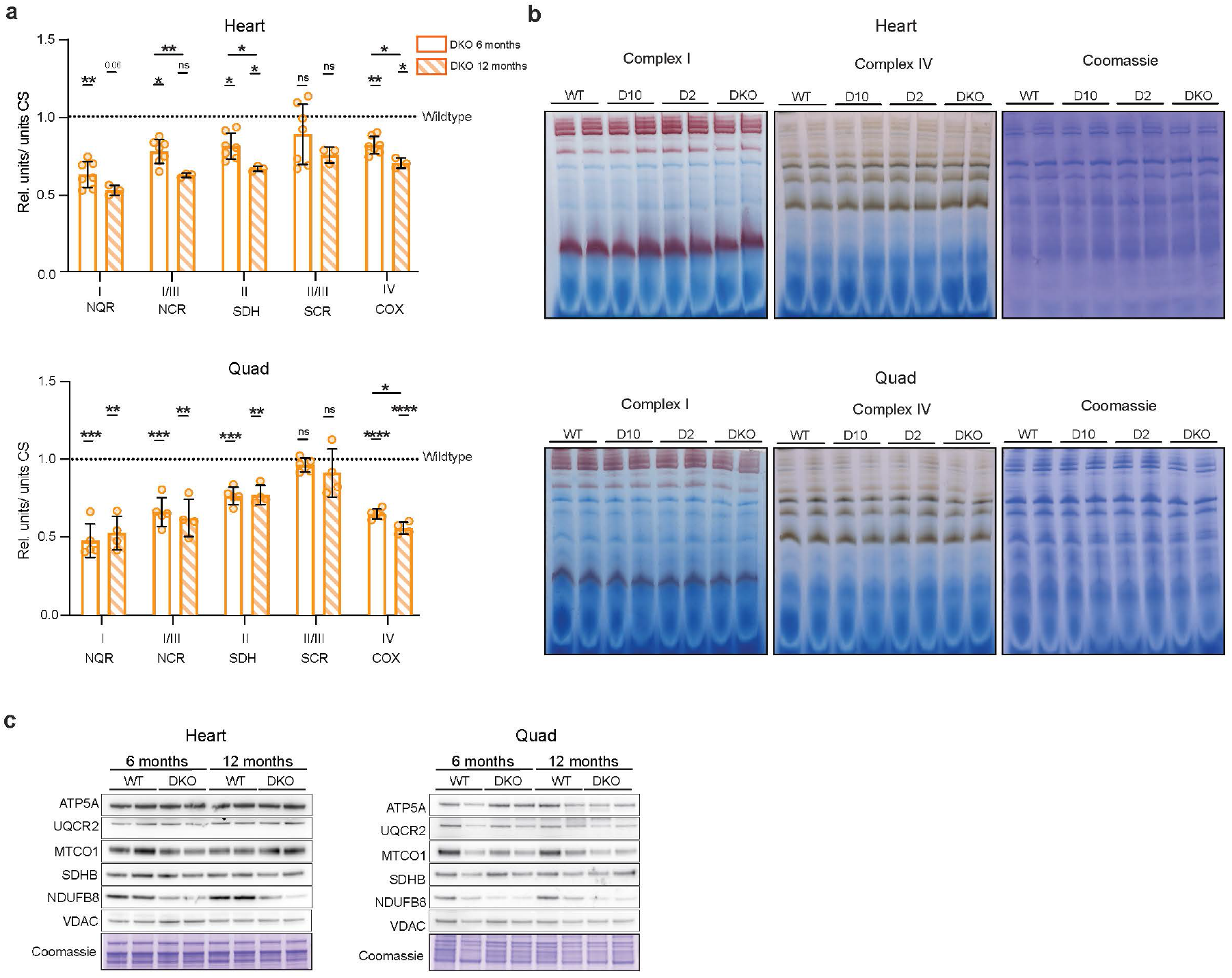
Loss of CHCHD2 and CHCHD10 compromises OXPHOS capacity in heart and skeletal muscle. **a,** Respiratory chain activities of CI, CI+CII, CII, CII+CIII, and CIV were measured spectrophotometrically in the heart and quadriceps of WT and DKO mice at 6 and 12 months of age. Enzyme activities were normalized to citrate synthase activity and expressed relative to WT controls (dotted line). Data are presented as means ± SD (n> 3 per genotype and timepoint). *p < 0.05; **p < 0.01; ***p < 0.001; ****p < 0.0001; ns, not significant. **b,** In-gel activities of complexes I and IV in digitonin-solubilized mitochondria isolated from heart and quadriceps of WT, D2 KO, D10 KO, and DKO mice at 6 months of age. Coomassie staining was used as a loading control. **c,** Western blot of steady-state OXPHOS subunits in mitochondria isolated from WT and DKO heart and quadriceps at 6 and 12 months of age. VDAC and Coomassie staining were used as loading controls.

To account for metabolic differences among skeletal muscle groups, we also analyzed the soleus, a slow-twitch, OXPHOS-dependent muscle^35^. In-gel activity assays of mitochondria isolated from 6- and 12-month-old soleus demonstrated a reduction in complex I similar to that observed in quadriceps (Supplementary Fig.2b).

Consistent with these findings, steady-state levels of OXPHOS subunits were reduced in mitochondrial extracts from DKO tissues at both 6 and 12 months of age, with a pronounced reduction in the complex I NDUFB8 evident at 6 months (Fig. 2c; Supplementary Fig. 2c-e). To assess whether the described defects were caused by alterations in mtDNA copy number, as previously reported in heart and skeletal muscle from CHCHD10 mutant mice^36^, we analyzed mtDNA levels. No changes were detected in 6-month-old DKO mice, suggesting that OXPHOS deficiency is not caused by impaired mtDNA homeostasis (Supplementary Fig. 2f).

### Tissue-specific proteomic responses induced by CHCHD2-CHCHD10 deficiency

To define molecular pathways affected by CHCHD2 and CHCHD10 loss, we performed label-free quantitative proteomics on whole-tissue extracts from the hearts and quadriceps of D2 KO, D10 KO, and DKO male and female mice at 6 months of age. Approximately 7,500 proteins were quantified in the heart and 5,500 in the quadriceps. Principal component analysis (PCA) revealed a separation of DKO heart samples from other genotypes, whereas quadriceps samples showed no clear clustering (Supplementary Fig. 3a, b). Consistently, more than 300 proteins were significantly altered in DKO hearts, while only five reached statistical significance in skeletal muscle (Fig. 3a). No significant differences were detected between D10 KO, D2 KO and controls in either tissue.

**Fig. 3:**
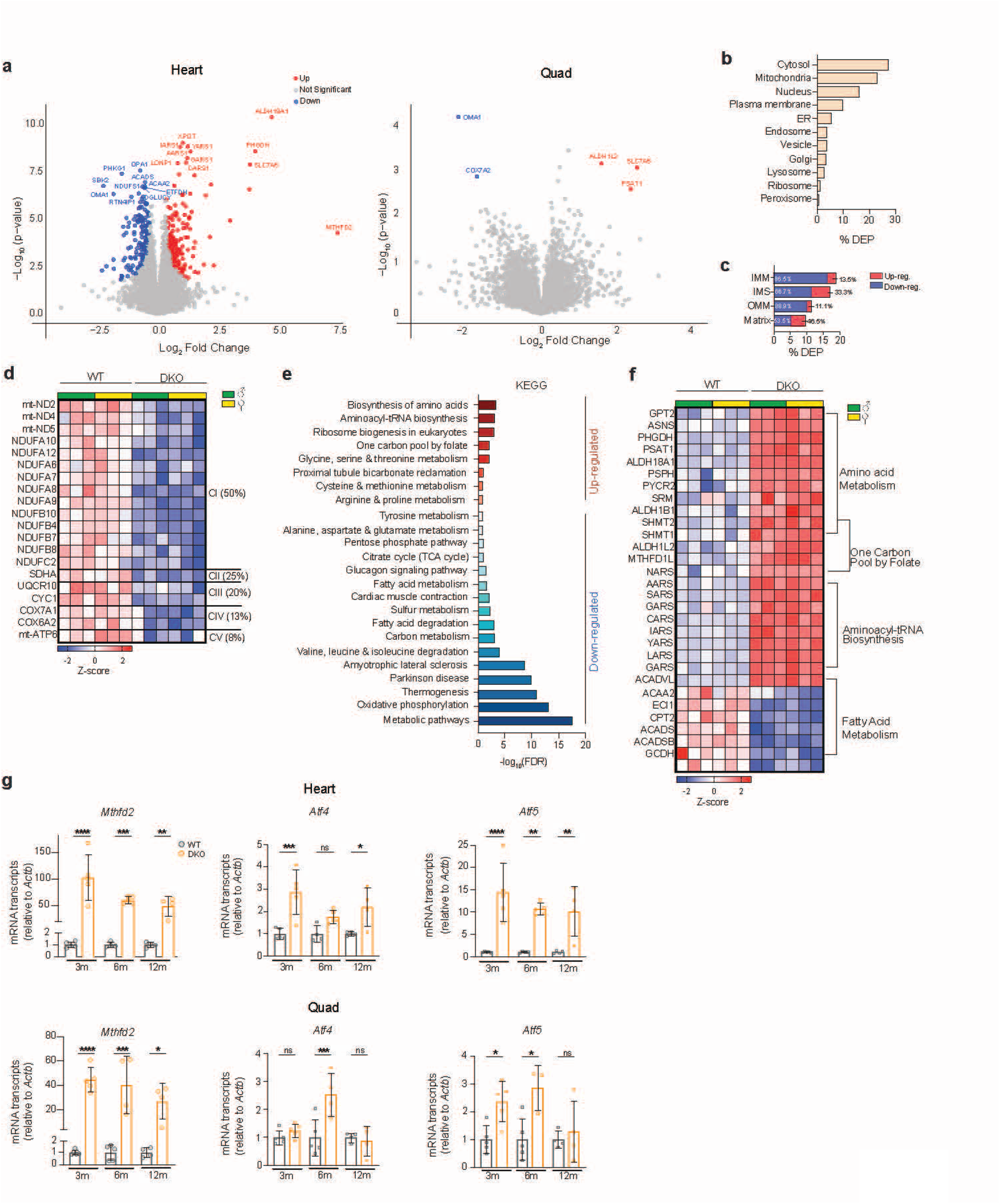
Proteomics reveals distinct tissue-specific molecular signatures in DKO mice. **a,** Volcano plots (log_2_ fold change versus - log_10_(p value)) of heart and quadriceps proteomics from 6-month-old DKO compared to WT mice. Significantly upregulated proteins are denoted in red, downregulated proteins in blue, and unchanged in grey. Each dot represents a single protein. Proteins with FDR <0.05 were considered significant. **b,** Distribution of differentially expressed proteins (DEPs) of DKO relative to WT by cell compartment based on GO cellular component annotations from heart proteomics. **c,** Percentage of DEPs between WT and DKO mitochondrial compartments based on MitoCarta 3.0 localization, ensuring compartment-specific effects are relative to the known distribution of mitochondrial proteins. Bar segments indicate the percentage up- and downregulated proteins relative to their contribution within each compartment. **d,** Heatmap of heart proteomics showing significantly changed OXPHOS proteins with percentage DEPs per OXPHOS complex. **e,** KEGG pathway enrichment analysis from heart proteomics at 6 months of age from DKO mice. Pathways denoted in red are enriched (upregulated), while blue are depleted (down-regulated). **f,** Heatmap of heart proteomics showing significantly changed proteins identified in KEGG pathway analysis. Each column represents an individual mouse; protein abundance is displayed as Z-scores. **g,** mRNA expression of *Mthfd2*, *Atf4* and *Atf5* relative to *Actb* from heart and quadriceps tissue of 3,6 and 12-month-old WT and DKO mice. Data presented as means ± SD (n > 4 per genotype and timepoint). *p < 0.05; **p < 0.01; ***p < 0.001; ****p < 0.0001; ns, not significant.

Compartment-specific analysis of cardiac differentially expressed proteins (DEPs) revealed that proteomic alterations were not confined to mitochondria but also extended to the cytosol and nucleus. Within mitochondria, normalization of DEPs to MitoCarta 3.0 compartment annotations showed that the largest fraction localized to the IMS and IMM, where proteins were predominantly downregulated (Fig. 3c). This highlights pronounced perturbations in the mitochondrial compartments occupied by, or immediately adjacent to, CHCHD2 and CHCHD10.

KEGG pathway enrichment analysis in the heart identified that downregulated pathways were dominated by OXPHOS, with approximately 50% of complex I subunits significantly decreased (Fig. 3d), consistent with complex I deficiency observed in these mice (Fig. 2). Additional reductions in fatty acid metabolism, the tricarboxylic acid cycle (TCA), and the pentose phosphate pathway further indicate broad suppression of mitochondrial oxidative and catabolic processes (Fig. 3e,f). In parallel, upregulated proteins revealed coordinated metabolic reprogramming characterized by the induction of amino acid biosynthetic pathways, including glycine, serine, and threonine metabolism, aminoacyl-tRNA biosynthesis, and ribosome biogenesis (Fig. 3e,f).

Several enzymes involved in one-carbon metabolism, including MTHFD2, were upregulated, suggesting the activation of mitochondrial stress-response pathways (Fig. 3a). In line with previous studies linking CHCHD10 mutations to ISR signaling^24,37,38^, enrichment analysis revealed a significant increase in ATF4 downstream targets, including PHGDH, PSAT1, and ASNS (Supplementary Fig. 3d). Notably, the stress protease OMA1 was markedly reduced, reflecting its activation and subsequent autocatalytic degradation^9^. This interpretation is supported by the near-complete cleavage of its substrate OPA1, as indicated by the loss of long OPA1 isoforms and the accumulation of short forms (Supplementary Fig. 3e). In addition, other proteins involved in mitochondrial dynamics, including MFN1, MFN2, and DNM1L, were reduced, whereas cristae-stabilizing proteins such as CHCHD6 and GHITM (TMBIM5) were upregulated (Supplementary Fig. 3f).

In skeletal muscle, despite minimal proteomic alterations under stringent statistical thresholds, OMA1 emerged as the most significantly downregulated protein (Fig. 3a), accompanied by similar changes in OPA1 processing (Supplementary Fig. 3e). Applying ANOVA-based criteria revealed additional modest changes, including reductions in OXPHOS-related proteins, predominantly complex I subunits, and mild induction of amino acid biosynthesis, one-carbon metabolism, and ATF4-linked ISR components. Although similar to those observed in the heart, these changes were substantially weaker (Supplementary Fig. 3g).

To characterize the temporal molecular dynamics, we quantified transcript levels of ISR markers at 3, 6, and 12 months of age (Fig. 3g). In DKO hearts, *Mthfd2*, *Atf4*, and *Atf5* were robustly induced by 3 months and remained elevated at later time points, although their expression was partially attenuated over time. In quadriceps, ISR-associated transcripts were induced at lower amplitude; *Mthfd2* showed the most consistent increase, whereas *Atf4* and *Atf5* exhibited modest and transient induction, consistent with a limited transcriptional response that did not translate into extensive proteomic changes.

Taken together, these data demonstrate that loss of CHCHD2 and CHCHD10 elicits a differential response across tissues. Although both heart and skeletal muscle exhibit marked respiratory chain defects, the heart undergoes extensive proteomic adaptive remodeling characterized by suppression of mitochondrial metabolism and early activation of ISR-linked biosynthetic programs, whereas skeletal muscle shows only minor alterations.

### Loss of the CHCHD2-CHCHD10 complex disrupts mitochondrial morphology and lipid composition

As CHCHD2 and CHCHD10 have been implicated in the maintenance of mitochondrial ultrastructure^39^, we performed transmission electron microscopy (TEM) on heart and quadriceps tissues from 6-month-old mice to assess tissue morphology and mitochondrial architecture. We used an automated segmentation approach to identify healthy and disrupted mitochondria, enabling the quantification of mitochondrial number, area, and aspect ratio (Fig.4a)^40^. In skeletal muscle, mitochondrial number was unchanged, whereas mitochondrial area was significantly increased across muscle sections (Fig. 4b,c), indicating altered mitochondrial morphology and consistent with increased swelling. In addition, skeletal muscle displayed an increased abundance of tubular aggregates (Fig. 4b; Supplementary Fig. 4a), corresponding to sarcoplasmic reticulum accumulations, which are typically associated with age-related muscle disorders^41^. In contrast, the heart exhibited a striking accumulation of markedly enlarged mitochondria with severely disrupted cristae (Fig.4b; Supplementary Fig.4b). These aberrant organelles, previously described as megamitochondria^42^, have often been associated with altered mitochondrial dynamics and reported in diverse pathological contexts^43–45^. Importantly, the majority of mitochondria (90%) retained structural integrity but displayed increased aspect ratio, consistent with elongation of the mitochondrial network (Fig. 4b,d; Supplementary Fig. 4e). In D10 KO hearts, megamitochondria were not detected, although mitochondrial size was mildly increased (Supplementary Fig. 4c-e).

**Fig. 4:**
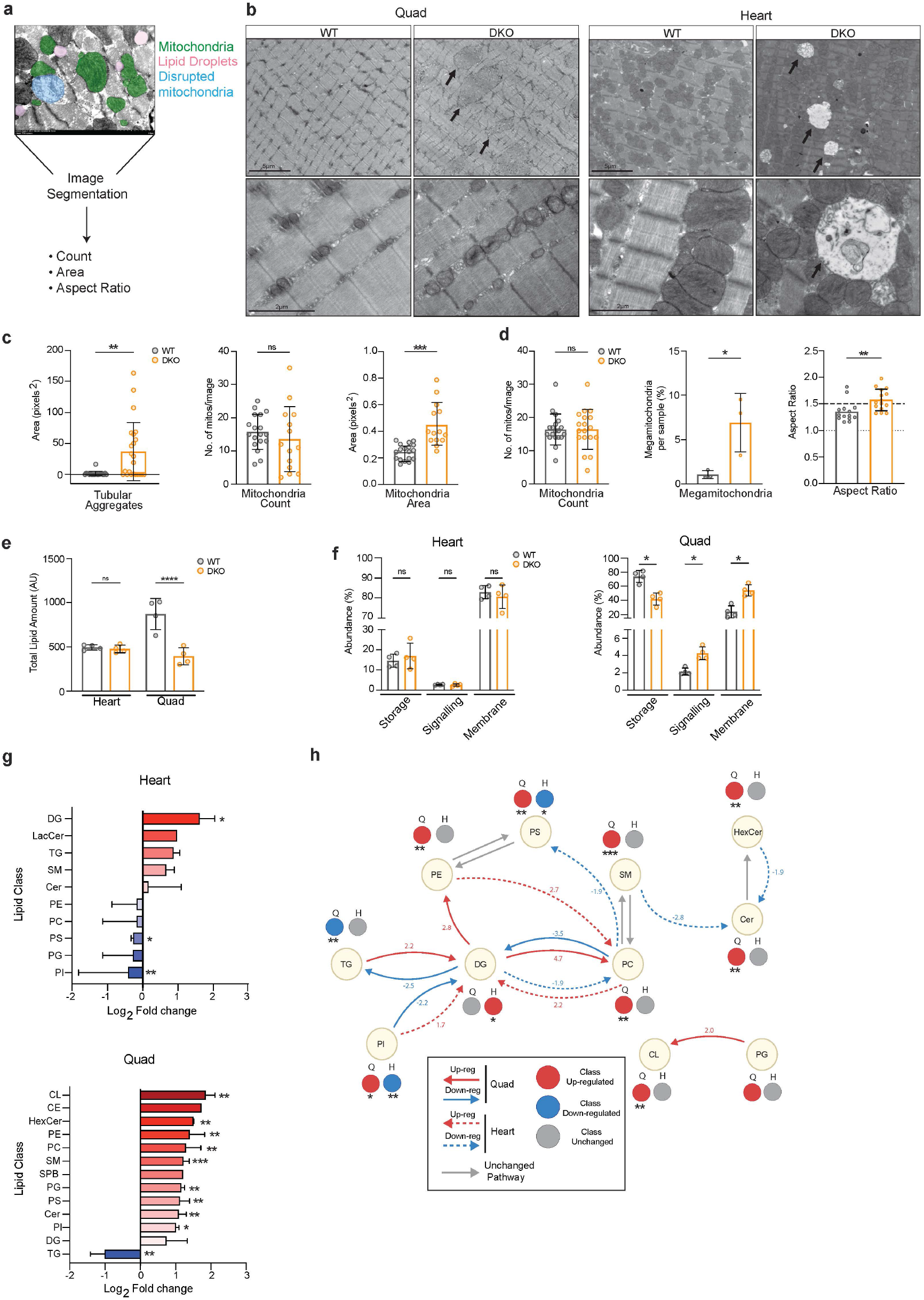
CHCHD2 and CHCHD10 ablation impacts mitochondrial morphology and lipid homeostasis in heart and skeletal muscle. **a,** Schematic illustrating the workflow for mitochondrial transmission electron microscopy (TEM) segmentation for downstream analyses. **b,** Representative TEM images at low and high magnification from quadriceps and heart of WT and DKO mice at 6 months of age. Scale bars 5µm (low magnification) and 2µm (high magnification). Quantification of mitochondrial morphology from TEM images acquired at 10,000X from **c,** quadriceps and **d,** heart (n= 3 mice with 5 images per replicate). **e,** Total lipid abundance measured by mass spectrometry in heart and quadriceps from 6-month-old male mice. Data is presented as means ± SD (n= 4). *p < 0.05; **p < 0.01; ***p < 0.001; ****p < 0.0001; ns, not significant. **f,** Relative abundance of lipid species grouped by biological function. Data presented as means ± SD (n= 4). *p < 0.05; **p < 0.01; ***p < 0.001.**g,** log_2_ fold change of lipid classes from heart and quadriceps lipidomics at 6 months of age. **h,** Network representation of lipid class alterations in the quadriceps and heart of 6-month-old DKO mice relative to WT. Nodes represent lipid classes and arrows indicate metabolic connections. Red and blue arrows denote significantly increased or decreased pathways, respectively (|log_2_ fold change| ≥ 0.5), with solid lines indicating skeletal muscle and dashed lines indicating heart; gray arrows indicate unchanged pathways. Pathway arrows are annotated with mean log_2_ fold changes of pathway scores. Lipid abbreviations are described in table 3.

We previously reported that CHCHD2 loss results in an accumulation of lipid droplet-like structures in the heart and subtle alterations in lipid homeostasis in affected brain regions^32^. Quantification of these structures revealed a modest, non-significant increase in D10 KO hearts, whereas no changes were observed in DKO mice (Fig. 4a; Supplementary Fig. 4e). To quantitatively assess lipid composition, we performed targeted lipidomic analysis of heart and quadriceps from 6-month-old mice. Total lipid content in skeletal muscle of DKO mice was substantially reduced (Fig. 4e,f), primarily due to depletion of storage lipids, particularly triglycerides (TGs) (Fig. 4g; Supplementary Fig. 4j). In parallel, several membrane-associated lipids, including phosphatidylinositol (PI), phosphatidylcholine (PC), and phosphatidylethanolamine (PE), were increased. Pathway analysis suggests a shift away from lipid storage toward mobilization, with preferential conversion of TGs into diacylglycerides (DG) intermediates and subsequent incorporation into structural phospholipids (Fig. 4h). The concomitant decrease in TGs with increased DG and phospholipids is consistent with enhanced lipid remodeling and redistribution, although altered lipid synthesis cannot be excluded. Cardiolipin (CL) species were also increased across multiple acyl-chain compositions, indicating perturbations of the IMM lipid homeostasis. However, the PC: PE ratio was unchanged between WT and DKO, suggesting no major alterations in membrane rigidity (Supplementary Fig.4k).

By contrast, the heart displayed no significant change in total lipid abundance nor in the relative distribution of major lipid classes (Fig. 4e,f). Nonetheless, analysis of individual species and pathway relationships revealed a distinct pattern. The DG-PC axis appeared to shift in the opposite direction compared to skeletal muscle (Fig. 4h; Supplementary Fig. 4j,l), yet overall membrane lipid composition remained unaffected. Consistent with this, CL species were not broadly altered, despite the presence of megamitochondria. The preserved lipid content is concomitant with downregulation of fatty acid metabolism pathways (Fig. 3f), suggesting that reduced lipid utilization may contribute to the maintenance of membrane lipid pools in the heart. Similarly, targeted lipidomic analysis of 12-month-old CHCHD10 KO hearts revealed minimal alterations, with only four lipid species significantly affected (Supplementary Fig. 4i). This contrasts with CHCHD10 S55L mutant hearts, which display pronounced accumulation of phospholipid and CL species^46^.

Overall, these findings highlight tissue-specific differences in lipid handling following CHCHD2 and CHCHD10 loss. Skeletal muscle exhibits redistribution of lipid stores, consistent with ongoing lipid mobilization. In contrast, the heart maintains a relatively stable lipid composition, likely reflecting the predominance of structurally intact mitochondria.

### CHCHD2 and CHCHD10 loss does not impair cardiac contractility and diverges from the CHCHD10 S55L phenotype

Given that knock-in mice carrying the CHCHD10 S59L or G58R variants develop a progressive mitochondrial cardiomyopathy leading to premature lethality^47,48^, we next asked whether the loss of CHCHD2 and CHCHD10 compromises cardiac function.

We focused on cardiac-associated signatures within our proteomic dataset and identified a reduction in proteins involved in cardiac contractility (e.g., MYH7 and TNNT2) and calcium handling (e.g., ATP2A2/SERCA2 and RYR2), accompanied by an increase in proteins associated with structural remodeling (e.g., TAGLN and POSTN) (Fig. 5a). Heart weight remained unchanged across genotypes at both 6 and 12 months of age, with only a minor reduction observed in male CHCHD2 KO mice (Fig. 5b; Supplementary Fig. 5a). Consistently, transcript levels of cardiomyopathy-associated markers, including *Nppa* (encoding atrial natriuretic peptide, ANP) and *Nppb* (encoding B-type natriuretic peptide, BNP), were not elevated (Supplementary Fig. 5b).

**Fig. 5:**
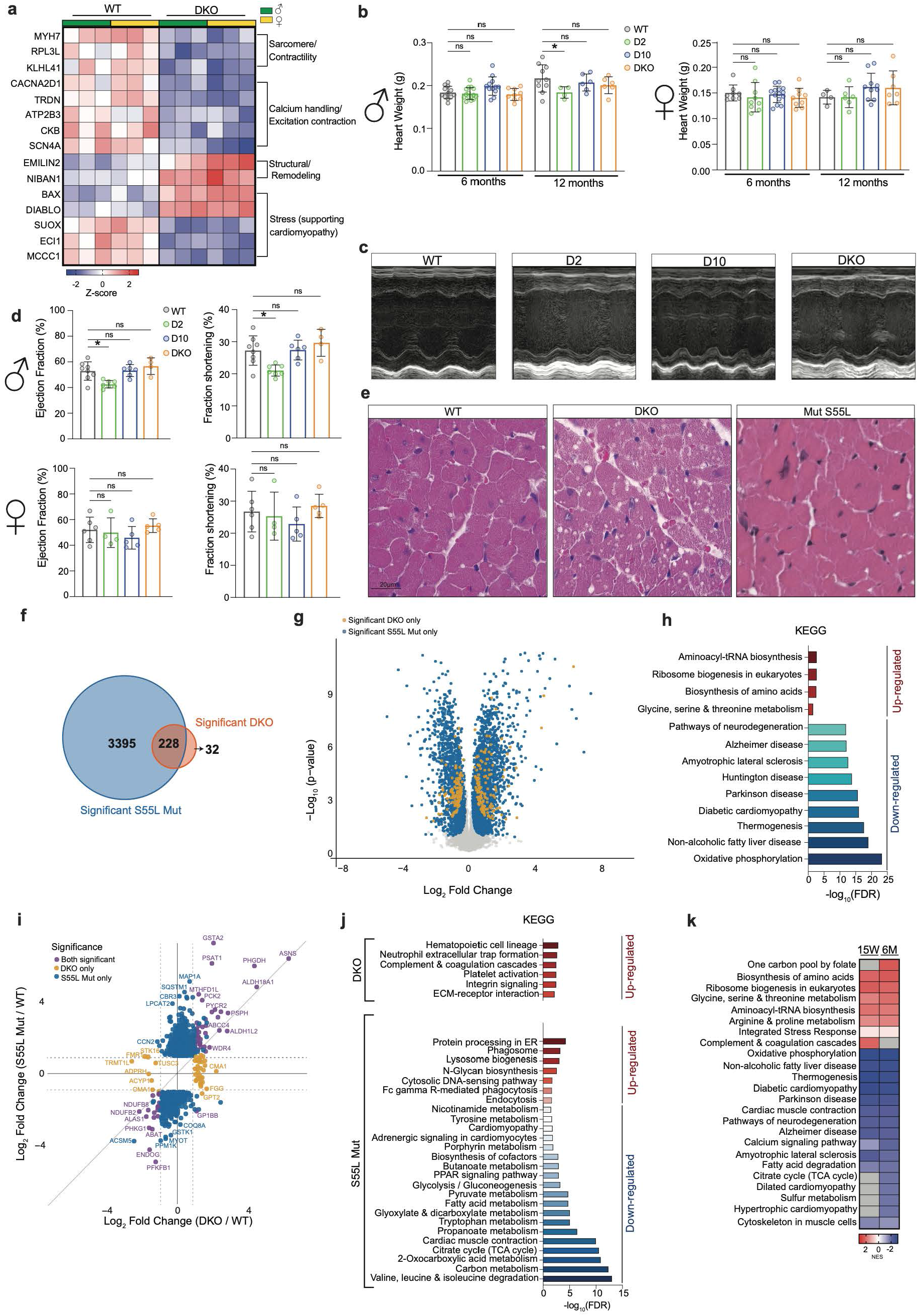
DKO mice preserve cardiac function in adulthood and do not phenocopy CHCHD10 S55L mutant pathology. **a,** Heatmap showing significantly changed cardiac-specific proteomic hits of WT and DKO mice at 6 months of age. Each column represents an individual mouse; protein abundance is displayed as Z-scores. **b,** Heart weight of WT, D2 KO, D10 KO and DKO male and female mice at 6 and 12 months of age. **C,** Representative echocardiography M-mode images of WT, D2 KO, D10 KO and DKO mice. **d,** Ejection fraction and fraction shortening measured from echocardiography of 6-8 month old WT, D2 KO, D10 KO and DKO male and female mice. Data are presented as means ± SD (n> 4 per genotype). *p <0.05, ns not significant. **e,** Representative histological hematoxylin and eosin staining of the left ventricle of the heart from WT, DKO and S55L mutant mice. Scale bar corresponds to 20µm. n> 3 animals and n> 4 technical replicates. **f,** Venn diagram showing overlap of significantly changed proteins (FDR < 0.05) from DKO and S55L mutant heart proteomics. **g,** Volcano plot (log₂ fold change versus −log₁₀(P value)) showing proteome overlap of DKO and S55L mutant heart at 3 months of age. Orange dots denote significantly changed proteins for the DKO, while blue highlights significant proteins unique to the S55L mutant. Gray dots are non-significant proteins. Proteins with FDR <0.05 were considered significant. **h,** KEGG pathway enrichment analysis from significant proteins from hearts at 3 months of age, common to both DKO and S55L mutant mice. Pathways denoted in red are enriched (upregulated), while blue are depleted (down-regulated). **I,** Log₂ fold change of DKO versus log₂ fold change of S55L mutant heart at 3 months, showing only significant (|Log₂ fold change| ≥ 1, FDR <0.05) proteins common to both mouse models (purple), unique to the DKO (orange), and the S55L mutant (blue). **j,** KEGG pathway enrichment analysis from significant proteins from hearts at 3 months of age, unique to DKO and S55L mutant mice. Pathways denoted in red are enriched (upregulated), while blue are depleted (down-regulated). **k,** Heatmap of normalized enrichment scores from KEGG pathway analysis comparing DKO hearts at 3 months and 6 months of age. Gray squares indicate non-significant pathways.

We next assessed cardiac performance by echocardiography between 6 and 9 months of age, when OXPHOS capacity was already substantially reduced. Despite the bioenergetic defect, DKO mice of both sexes, as well as CHCHD10 KO mice, retained normal heart contractility, with no evidence of impaired systolic function. In contrast, male CHCHD2 KO mice displayed a mild reduction in systolic performance, reflected by decreased ejection fraction and fractional shortening (Fig. 5c,d). Concomitantly, left ventricular internal diameter increased, whereas anterior wall thickness was reduced during both systole and diastole (Supplementary Fig. 5a), indicating that CHCHD2 ablation alone promotes a mild dilated cardiomyopathy. To investigate the mechanisms that preserve cardiac contractility in the DKO heart, we compared the DKO model with the pathogenic CHCHD10 S55L mutant, which develops with fatal cardiomyopathy^24,38^. Although S55L mutant hearts display pronounced ventricular wall thickening and reduced lumen consistent with hypertrophic cardiomyopathic remodeling^24^, cardiomyocyte architecture appeared normal in hematoxylin and eosin (H&E)-stained sections. In contrast, DKO hearts exhibited prominent intracellular accumulations consistent with vacuole-like structures and/or enlarged mitochondria (Fig. 5e), which were also evident in toluidine blue-stained semithin sections (Supplementary Fig. 5c,d).

To define early molecular determinants of divergent cardiac outcomes of DKO and S55L mutant mice, we performed proteomic analysis of heart tissue from 3-month-old animals. PCA revealed clear separation of both DKO and S55L mutant samples from their respective controls (Supplementary Fig. 5f). While the DKO exhibited approximately 250 significantly altered proteins, the S55L mutant showed a markedly broader proteomic response, with over 3,500 DEPs (Fig. 5f,g; Supplementary Fig. 5g).

Notably, ∼88% of proteins altered in the DKO were also differentially expressed in the S55L mutant (Fig. 5f,g). Enrichment analysis of commonly altered proteins identified reduced OXPHOS capacity and suppression of pathways associated with age-related human diseases (Fig. 5h, i). In parallel, aminoacyl-tRNA biosynthesis and amino acid metabolism were elevated, indicating activation of a shared stress-response program.

To further resolve model-specific differences, we directly compared fold changes between DKO and S55L hearts using an FDR < 0.05 and an additional log₂FC cutoff of 1. This approach distinguished proteins uniquely altered in each model and highlighted substantial differences in the magnitude of common molecular programs (Fig. 5i). Notably, DKO hearts showed no significantly enriched downregulated pathways and only a small number of selectively decreased proteins (Table 1). Instead, they exhibited selective enrichment of proteins associated with extracellular matrix interactions, platelet activation, and complement/coagulation pathways (Fig. 5j), suggesting engagement of non-cardiomyocyte remodeling programs that were not detected in S55L mutant hearts. Among proteins significantly altered in both models, a subset exhibited opposite expression patterns (Supplementary Fig. 5h), including mitochondrial gene expression factors such as POLRMT and TFB1M, indicating activation of compensatory mechanisms that were suppressed in S55L mutant hearts.

**Table 1:** Significantly downregulated proteins in 3-month-old DKO heart, unchanged in S55L Mutant.

| Protein | Log<sub>2</sub>FC | P-value |
| --- | --- | --- |
| NDUFA7 | -1,77262 | 0,012964 |
| SBK2 | -1,75301 | 0,006968 |
| STK16 | -1,63522 | 0,027909 |
| SCN4a | -1,4675 | 0,009528 |
| ASB5 | -0,99809 | 0,049672 |
| DACT3 | -0,97405 | 0,016199 |
| ARID2 | -0,95964 | 0,046152 |
| COX6C | -0,86622 | 0,045498 |
| COX19 | -0,81062 | 0,012109 |
| COA7 | -0,67216 | 0,020514 |
| SLC25A46 | -0,62607 | 0,019275 |
| CKB | -0,62215 | 0,029265 |
| NENF | -0,58972 | 0,04998 |
| PLIN4 | -0,5843 | 0,05000 |

The S55L mutant hearts presented widespread pathway alterations not detected in the DKO, including activation of proteostasis, intracellular trafficking and lysosomal pathways (Fig. 5j). Mutant hearts also showed broad suppression of central metabolism, affecting glycolysis, pyruvate oxidation, fatty acid utilisation, the TCA cycle, and branched-chain amino acid degradation. Direct comparison of the two proteomes further revealed a prominent immune-response signature unique to the S55L model, in line with previous evidence linking overactive immune activation to disease severity^38^. Moreover, although both models activated the ISR, the response was substantially stronger in S55L hearts (Supplementary Fig. 5i). We next examined the temporal evolution of the proteomic response in DKO hearts by comparing mice at 3 and 6 months of age. A substantial fraction of the proteomic alterations detected at 3 months persisted at 6 months, indicating the establishment of an early and sustained adaptive response. Pathway enrichment analysis further revealed only minor progressive molecular changes. While OXPHOS-associated proteins slightly declined further, amino acid metabolism remained elevated. Few responses emerged at 6 months, most notably the activation of one-carbon metabolism. In fact, MTHFD2 was not detected at the protein level at 3 months but was strongly induced by 6 months. Likewise, proteomic signatures associated with cardiac contractility and structural adaptation became apparent only at 6 months, whereas the early enrichment of complement and coagulation pathways was no longer observed (Fig. 5k).

Overall, despite progressive bioenergetic defects, mice lacking CHCHD2 and CHCHD10 maintain cardiac function through a mild and adaptive proteomic response. By contrast, the CHCHD10 S55L mutation triggers extensive detrimental metabolic, proteostatic, and immune dysregulation that ultimately culminates in fatal cardiomyopathy.

### CHCHD2-CHCHD10 deficiency impairs muscle satellite cell function and myogenic progression

To investigate the basis of the skeletal muscle phenotype observed in DKO mice, characterized by reduced lean mass and muscle weakness, we performed morphometric analysis of quadriceps muscles from 6-month-old mice using laminin immunostaining. DKO muscles exhibited a significant reduction in myofiber cross-sectional area (CSA), reflecting a shift in fiber size distribution (Fig. 6a,c), which has previously been reported in mitochondrial mouse models of myopathy^49,50^. Importantly, no differences in myofiber size were detected in S55L mutant muscle at 4 months of age (Fig. 6b,d). Consistent with this finding, S55L mice also showed no reduction in muscle mass or strength at this age, whereas both phenotypes became apparent from 6-7 months of age^24^. Centrally located myonuclei, a marker of ongoing degeneration-regeneration cycles^51^, were not affected in DKO muscle (Fig. 6e). However, the expression of the myogenic regulators *Pax7* and *Myod1* was reduced at 6 months (Fig. 6f), suggesting impaired myogenic regenerative capacity.

**Fig. 6:**
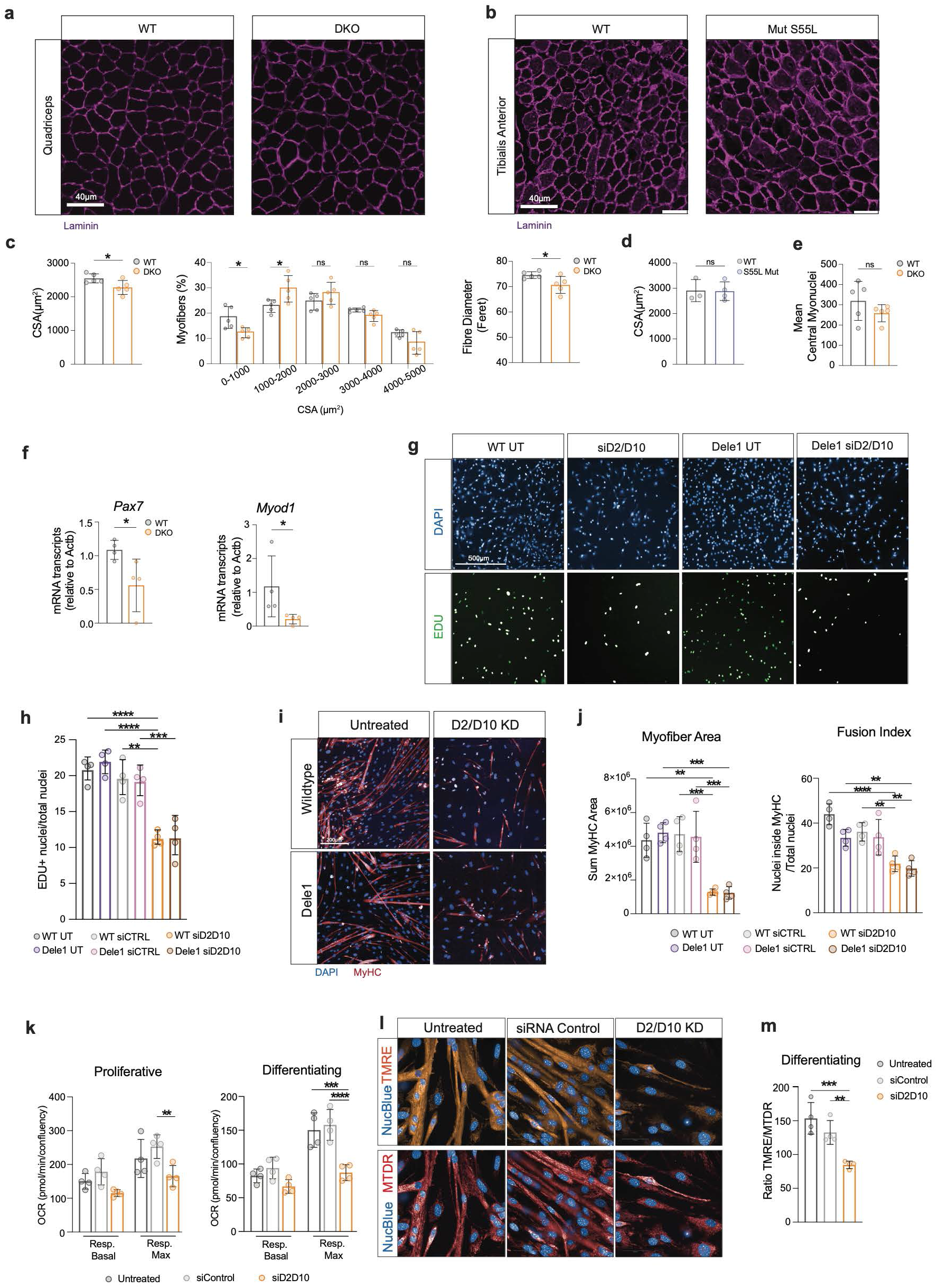
CHCHD2 and CHCHD10 deficiency impairs muscle fiber size and satellite cell function. Representative images of laminin stained muscle cross sections from **a,** quadriceps from DKO mice at 6 months and **b,** tibialis anterior CHCHD10 S55L mutant muscle for morphometric analysis of muscle fibers. n>3 animals, scale bar = 40µm. **c,** Quantification of cross-sectional area and diameter (feret) of DKO muscle fibers. Data are presented as means ± SD (n = 5), *p < 0.05. **d,** Quantification of cross-sectional area of S55L mutant muscle fibers. Data are presented as means ± SD (n > 3), ns not significant. **e,** Mean central myonuclei quantification from DKO quadriceps at 6 months of age. Data are presented as means ± SD (n = 5), ns not significant.**f,** mRNA transcript levels of *Pax7* and *Myod1* relative to *Actb* from 6-month-old WT and DKO mice. Data presented as means ± SD (n = 4) *p < 0.05. **g,** Representative images of DAPI and Edu labeled primary satellite cells of WT, *Dele1* KO and S55L mutant mice, with and without siD2/D10. Scale bar: 500µm. **h,** Quantification of Edu+ nuclei of labeled primary satellite cells of WT and *Dele1* KO mice, with and without siD2/D10. Data are presented as means ± SD (n = 4) **p < 0.01; ***p < 0.001; ****p < 0.0001. **i,** Representative images of WT and *Dele1* KO,sicontrol and siD2/D10 treated differentiating satellite cells after 48H, stained with DAPI and MyHC. Scale bar: 200µm. **j** Quantification of myofiber area and fusion index after 48H of differentiation of WT and Dele1 KO satellite cells treated with siD2/D10. Data presented as means ± SD (n = 4) **p < 0.01; ***p < 0.001; ****p < 0.0001. **k,** Basal and maximum respiration of proliferative and differentiating WT satellite cells treated with siD2/D10. Data presented as means ± SD (n = 4) **p < 0.01; ***p < 0.001; ****p < 0.0001. **i,** Representative images of TMRE and MTDR-stained differentiating WT satellite cells treated with siD2/D10. Scale bar: 50µm. **m,** Quantification of the ratio of TMRE to MTDR in differentiating WT satellite cells treated with siD2/D10. Data are presented as means ± SD (n = 4) **p < 0.01; ***p < 0.001.

Given the importance of mitochondria in maintaining the regenerative potential of adult muscle stem cells ^52,53^, also known as satellite cells, we investigated whether loss of CHCHD2 and CHCHD10 directly affects satellite cell function. To this end, primary satellite cells isolated from adult wild-type mice were depleted of *Chchd2* and *Chchd10* by siRNA (Supplementary Fig. 6a). Combined knockdown markedly reduced cell proliferation, as assessed by EdU incorporation (Fig. 6g,h; Supplementary Fig. 6b-d). This defect was associated with impaired myogenic differentiation, evidenced by reduced myotube formation, smaller myotube area, and a lower fusion index at both 24 and 48 h (Fig. 6i, j; Supplementary Fig. 6e, f). In contrast, satellite cells isolated from S55L mutant mice displayed normal proliferative and differentiation capacities (Supplementary Fig. 6c, d), indicating that the muscle phenotype associated with this pathogenic mutation arises through mechanisms distinct from satellite cell dysfunction.

As OMA1 activation and the mitochondrial ISR were among the few molecular pathways induced in DKO skeletal muscle, we next examined whether this pathway mediates the satellite cell defects. We therefore isolated satellite cells from adult *Dele1* knockout mice. Loss of DELE1 alone did not affect cell proliferation or differentiation, and combined depletion of CHCHD2-CHCHD10 in a DELE1-deficient background did not impact the observed defects (Fig. 6g-j), indicating that impaired myogenic cell functions occur independently of DELE1-mediated ISR signaling.

To determine whether these deficits instead arise from an intrinsic bioenergetic impairment, we assessed mitochondrial respiratory capacity in proliferating and differentiating satellite cells. Basal respiration was unchanged, whereas maximal respiratory capacity was selectively reduced during differentiation following CHCHD2-CHCHD10 depletion (Fig. 6k; Supplementary Fig. 6k, n). Consistently, mitochondrial membrane potential, assessed by TMRE staining, remained normal in proliferating cells but markedly decreased during differentiation (Fig. 6l, m; Supplementary Fig. 6j, k). Importantly, the mitochondrial membrane potential was normal in satellite cells carrying the CHCHD10 S55L mutation (Supplementary Fig. 6l, m).

Together, these findings demonstrate that loss of CHCHD2 and CHCHD10 impairs satellite cell proliferation and differentiation and is associated with reduced mitochondrial bioenergetic capacity during myogenesis. In support of this model, CHCHD10 expression is dynamically induced during myogenic progression, increasing markedly upon satellite cell activation and differentiation^54^. Notably, these defects were not recapitulated in satellite cells carrying the pathogenic CHCHD10 S55L mutation and occurred independently of DELE1-mediated ISR signaling.

## Discussion

Unlike core mitochondrial processes inherited from prokaryotic ancestors, IMS-resident proteins evolved in eukaryotes to support the dynamic flexibility and proteome surveillance required for tissue function^55,56^. Among these, CHCHD2 and CHCHD10 have attracted considerable attention due to their association with an expanding spectrum of human diseases^23^. Phylogenetic analyses indicate that *CHCHD2* and *CHCHD10* arose from a gene duplication event early in vertebrate evolution^57^. While vertebrates express both paralogs, invertebrates such as *D. melanogaster* and *C. elegans*, as well as unicellular eukaryotes like *S. cerevisiae*, harbor only a single *CHCHD2/10*-like ortholog, indicating that duplication was followed by functional specialization^30,58,59^. Consistent with partial redundancy, ablation of either CHCHD2 or CHCHD10 results in mild or negligible phenotypes in mice ^28,32,60^. In striking contrast, many disease-associated variants cause severe pathology through dominant gain-of-function mechanisms^24,38,61^. Certain mutant CHCHD2 and CHCHD10 proteins misfold and accumulate as large insoluble aggregates within the IMS, sequestering their endogenous counterparts and impairing the native complex function. This dual gain- and loss-of-function mechanism has confounded efforts to define the physiological role of the CHCHD2-CHCHD10 complex.

Here, we provide the first comprehensive characterization of whole-body *Chchd2/Chchd10* DKO mice across adulthood, with a focus on cardiac and skeletal muscle. At the biochemical level, DKO animals exhibit clear defects in OXPHOS capacity, most prominently affecting complex I in both tissues. Notably, these defects occur in the absence of major alterations in the assembly of individual respiratory complexes and instead are accompanied by substantial abnormalities in mitochondrial morphology, including swollen and structurally aberrant organelles. These results suggest that the CHCHD2-CHCHD10 complex contributes to the organization of the IMM, likely impacting the higher-order respiratory chain through effects on membrane lipid composition and proteome stability.

A striking feature of our findings is the pronounced tissue-specific response to CHCHD2 and CHCHD10 ablation. Despite comparable degrees of OXPHOS impairment, skeletal muscle and heart exhibit markedly distinct responses and molecular phenotypes. The heart of DKO mice underwent early proteomic remodeling while maintaining overall lipid homeostasis, even in the presence of pronounced accumulation of disrupted megamitochondria. In contrast to a previous report^26^, we found no evidence of impaired cardiac contractility by echocardiography. Interestingly, mild cardiomyopathic features were only detected in CHCHD2-deficient males, which may reflect insufficient activation of early adaptive stress responses that are otherwise triggered by the additional ablation of CHCHD10^32^. Consistent with this hypothesis, young DKO hearts displayed activation of multiple adaptive pathways, including the ISR, one-carbon metabolism, amino acid biosynthesis, and mitochondrial gene expression programs, which may help support cardiac performance into adulthood. However, late-onset cardiomyopathy cannot be excluded, as proteomic analyses revealed the gradual emergence of signatures associated with cardiac remodeling and contractile dysfunction over time.

In skeletal muscle, the response to CHCHD2 and CHCHD10 loss was profoundly different. Despite minor changes in the tissue proteome, DKO muscle exhibited extensive lipid remodeling, characterized by depletion of TG stores and an accumulation of membrane lipids, indicative of enhanced lipid mobilization. In mature myofibers, neutral lipid stores constitute an important energy reservoir that supports mitochondrial metabolism^62^. Thus, the pronounced shift away from lipid storage observed in DKO muscle may reflect a compensatory response to chronic mitochondrial dysfunction. However, the consequences of this remodeling may extend beyond mature myofibers. Alterations in lipid dynamics have been linked to defects in satellite cell proliferation and muscle regeneration^63^, raising the possibility that the observed lipid perturbations contribute to the impaired myogenic capacity observed following CHCHD2 and CHCHD10 depletion. Indeed, lipid remodeling in DKO was accompanied by reduced myofiber size and decreased expression of myogenic regulators, demonstrating that the CHCHD2-CHCHD10 complex is required for postnatal muscle growth and maintenance. This interpretation is further supported by the expression patterns of *Chchd2* and *Chchd10*. Both genes are expressed at low levels during embryonic myogenesis, whereas their expression increases substantially after birth and remains elevated throughout postnatal muscle maturation^54,64,65^.

The accumulation of tubular aggregates in DKO muscle provides additional evidence of impaired muscle homeostasis. In humans, tubular aggregates are observed in several myopathic conditions as well as during physiological aging^41,66^, and their presence in DKO muscle is therefore consistent with the emergence of an early muscle-aging phenotype. Notably, low *CHCHD10* expression has recently been proposed as a biomarker of early sarcopenia^67^, while muscle-specific ablation of CHCHD2 and CHCHD10 has been shown to recapitulate key features of muscle aging, including reduced ATP production ^68^. Consistent with these observations, our data indicate that the reduced OXPHOS capacity of mature DKO muscle may originate during myogenic differentiation, when impaired maximal respiratory capacity and mitochondrial membrane potential limit the metabolic flexibility required to sustain myogenesis. This defect appears independent of DELE1 signaling. Although DELE1 deletion has been shown to exacerbate the DKO phenotype^69^, the mitochondrial defects induced by CHCHD2-CHCHD10 deficiency in myogenic cells were not further aggravated by DELE1 loss, suggesting that mitochondrial stress-response pathways regulate disease progression in a tissue- and cell-type-specific manner.

Remarkably, the disease-associated CHCHD10 S55L mouse model diverges mechanistically from the DKO model. Unlike DKO mice, knock-in mice retain normal myogenic cell function, suggesting that the muscle weakness and atrophy observed in these animals arise from distal motor neuropathy, likely secondary to neuromuscular junction dysfunction ^24,25^.

Our findings demonstrate that DKO mice do not merely recapitulate the pathogenic CHCHD10 S59L mutation associated with mitochondrial disease in humans. Instead, they develop a unique phenotype that reveals differential vulnerability across tissues. The heart appears more capable of buffering metabolic and structural stress, whereas skeletal muscle is particularly vulnerable during growth and regeneration, leading to impaired myogenesis accompanied by altered lipid mobilization. Together, our study supports a model in which CHCHD2 and CHCHD10 form an IMS-localized regulatory complex that preserves mitochondrial integrity and supports muscle maintenance during postnatal growth and aging.

## Materials and Methods

### Ethical Statement

All procedures involving animals complied with ethical standards and followed all relevant European, national, and institutional guidelines. The Stockholm ethical committee approved all protocols and procedures (Ethical permit 18936-22), which were carried out in accordance with the guidelines set by the Federation of European Laboratory Animal Science Associations (FELASA). Animals were handled with a non-aversive handling approach. Animal work associated with the CHCHD10 S55L mutant mouse was carried out in France in accordance with guidelines set by the French Ministry of Research (Ethical permit APAFIS#32988-2021091417302194 v1).

### Mouse Models

All mice were bred on a C57BL/6N background. Mice were housed at 21°C, with 12h light-dark cycles and provided standard chow *ad libitum*. Whole-body homozygous *Chchd10* knockout mice were generated by the Karolinska Center for Transgene Technologies (Karolinska Institutet, Stockholm, Sweden) using CRISPR-Cas9. Briefly, C57BL/6NCrl zygotes (Charles River Laboratories) were microinjected with a pre-formed Cas9 ribonucleoprotein complex using two single guide RNAs (CRISPRevolution sgRNA EZ Kit, Synthego). Embryos were transferred into the oviducts of pseudopregnant surrogate mice via microsurgical transfer.

Thereby, a 138bp deletion spanning an exon-intron junction at the end of exon 2 of *Chchd10* was introduced, resulting in loss of CHCHD10 protein expression. Germline transmission of the deleted allele was confirmed by Sanger sequencing (KIGene, Karolinska Institutet, Stockholm, Sweden). Consequently, heterozygous *Chchd10* KO mice were intercrossed to generate homozygous knockouts (D10 KO). Genotyping was performed by PCR using genomic DNA isolated from ear biopsies following standard protocols.

NGL-Chchd10-g1 TGAGCCCACGGCTACGCCTG

NGL-Chchd10-g2 CTGACCAGTGCCTTCAGTGG

Whole body homozygous *Chchd2* knockout mice (D2 KO) were obtained as previously described^32^. CHCHD10 S55L heterozygous mutant mice were generated as previously described^25^.

To generate *Chchd2^-/-^* + *Chchd10^-/-^*double knockout mice (DKO), *Chchd2^-/-^* and *Chchd10^-/-^*were crossed, subsequently *Chchd2^-/-^ + Chchd10^+/-^* mice were intercrossed to generate both D2 KO as well as DKO mice. Additionally, *Chchd10^+/-^*mice were intercrossed to keep an experimental cohort of wildtype (WT) and Chchd10^-/-^ (D10 KO) mice. Breeding for both mouse lines was housed within the same room and timed together to ensure comparable conditions for all four genotypes. All analyses were performed blinded to genotype where possible. Group sizes and sex are indicated in figure legends.

### Behavioral Testing

Behavioral testing in WT, D2 KO, D10 KO, and DKO was conducted at 6 and 12 months of age for males and females separately. Before testing, all mice were acclimatised to the ventilated experimental room for at least 30 min. All tests were conducted at the same period of the light cycle (8 am- 6 pm).

The rotarod (Ugo Basile, Italy) was used to assess motor coordination and endurance. Mice were trained the day before testing in two 90-second sessions at a fixed rotation speed of 4 rpm. For the test, mice were placed on the rotarod in acceleration mode (4-40 rpm) for 5 minutes, and the latency to fall off the rod was recorded for analysis. Each animal had five trials with an inter-trial resting period of 5 minutes.

Muscle strength in all four limbs was determined using the grip strength apparatus (BioSeb, FL, USA). Mice were placed on a metal grid and encouraged to grip the grid with their forepaws and hind paws. By gently gripping and pulling the base of the tail, a horizontal force was applied. Peak force in grams (g) was gauged using a digital force gauge attached to the metal grid. Each animal was tested three times with 5 minutes of rest between measurements. The average of the three measurements was used for analysis.

### Body composition analysis

The fat and lean mass of animals was non-invasively determined 2 days before behavioral studies at 6 and 12 months of age using quantitative magnetic resonance imaging (MRI) (EchoMRI-100, EchoMRI).

### Echocardiography

Transthoracic echocardiography was performed using the VisualSonics Vevo 2100 system with an MS550D transducer. Before echocardiography, anaesthesia was conducted using 2% isoflurane for initial induction. After confirming the absence of pedal reflexes, anaesthesia was maintained at 1.2% - 1.3% isoflurane for the duration of acquisition. Body temperature and real-time heart rate were monitored with the Vevo physiology monitoring platform. Heart rate was maintained within the physiological range during acquisition. A parasternal short-axis M-mode scan at the mid-ventricular level was acquired to evaluate systolic function with papillary muscles as guidance. Left ventricular ejection fraction (EF), fractional shortening (FS), interventricular septum (IVS), left ventricular internal diameter (LVID), and left ventricular anterior wall thickness (LVAW) measured during systole and diastole were calculated based on the average of 5-7 cardiac cycles.

### Histology

#### Haematoxylin and Eosin staining

Heart tissues were stored in 4% paraformaldehyde (PFA) in phosphate-buffered saline (PBS) for 24 hours at 4°C, then transferred to 70% ethanol before processing. Samples were dehydrated using the Excelsior AS Tissue Processor (Epredia). Samples were embedded in warm paraffin wax and sectioned using the microtome (HM355s automated Microtome, Epredia). Slices of 3µm thickness were mounted onto glass slides (SuperFrost Plus, J1800AMNZ, Thermo Scientific). Heart sections were deparaffinized using xylene, followed by sequential rehydration and stained with hematoxylin and eosin according to standard laboratory protocols. Sections were imaged at 10X using the Axioscan Z.1 slide scanner (Zeiss). Histopathological evaluation was performed using QuPath software^70^.

#### Skeletal muscle morphometric analysis

Fresh frozen quadriceps and tibialis anterior embedded in OCT were transversally cryosectioned to 12-14µm and mounted on Polysine glass slides (631-0107 VWR). Slides were washed three times in PBS with 0.1% Triton X-100 for permeabilization, before blocking in 10% fetal bovine serum (FBS) in PBS for 1h in a humidified chamber. Sections were incubated in primary Laminin (see Table 2) antibody overnight at 4°C under humid conditions. Slides were washed in PBS three times to remove the primary antibody before incubation in the secondary antibody (Alexa Fluor 488, A11034, Thermo Fisher) in a dark, humidified chamber at room temperature for 2h. Slides were washed three times with PBS prior to DAPI staining and mounting using ProLong Gold Antifade mounting media.

**Table 2:** Primary Antibody List.

| Antigen | Source | Company | Catalog nr. | Dilution | Molecular weight (kDa) |
| --- | --- | --- | --- | --- | --- |
| CHCHD2 | Rat | Helmholtz Munich | CCHD 16D9 | Supernatant | 16 |
| CHCHD10 | Rabbit | Sigma | HPA003440 | (1:500) | 14 |
| CHCHD10 | Rabbit | ProteinTech | 19424-I-AP | (1:1000) | 15 |
| VDAC | Mouse | Abcam | MSA03/ab14734 | (1:2000) | 33 |
| MitoProfile Mouse | Mouse | Abcam | MS604-300 | (1:1000) | 55-48-40-30-20 |
| HSP60 | Mouse | Enzo Lifesciences | AB1-SPA-807-E | (1:2000) | 60 |
| HSC70 | Mouse | Santa Cruz | Sc-7298 | (1:2000) | 70 |
| OPA1 | Rabbit | Abcam | Ab42364 | (1:1000) | 90-100 |
| MTHFD2 | Rabbit | ProteinTech | 12270-1-AP | (1:1000) | 38 |
| ATP5A | Mouse | Abcam | Ab147448 | (1:1000) | 55 |
| MT-CO1 | Mouse | Invitrogen | Ab459600 | (1:1000) | 37 |
| CHCHD3 | Rabbit | ProteinTech | 25625-1-AP | (1:1000) | 26 |
| C1QBP (p32) | Rabbit | Sigma | HPA026483 | (1:1000) | 32 |
| LAMA1 | Rabbit | Sigma | L9393 | (1:500) | 400 |
| MyHC | Mouse | Developmental studies hybridoma bank | MF 20-c | (1:100-300) | 223 |

For morphometric fibre analysis and central myonuclei analysis, 3-4 whole muscle sections per mouse were imaged at 20X magnification using the Axioscan Z.1 slide scanner (Zeiss). Images were automatically analyzed using Fiji ImageJ (version: 2.16.0) and using the Plugin MuscleJ2^71^. Sections that did not pass the 40% artefact detection threshold or where the quality of fibre quantification was not uniform across sections were removed from subsequent analysis.

### Isolation of mitochondria from mouse tissue

Mice were euthanized by cervical dislocation or CO_2_, followed by decapitation, and several tissues were collected and rinsed in ice-cold PBS, including heart and skeletal muscle (quadriceps unless specified otherwise). Mitochondria were isolated with differential centrifugation, as previously described^32^. Crude mitochondria were snap-frozen in liquid nitrogen and stored at -80°C.

### Biochemical evaluation of respiratory chain function

Respiratory chain enzyme activity was measured from isolated mitochondria from heart and quadriceps as previously described^72^. Different enzyme activities were normalized to citrate synthase activity.

### Transmission electron microscopy

Small pieces of the proximal region (closest to the pelvis) of the left quadriceps and from the left ventricle of the myocardium were fixed immediately after dissection in 2% glutaraldehyde and 1% PFA in PBS and stored at 4°C. Samples were post-fixed, dehydrated, embedded and sectioned as described^32^. Ultrathin sections were contrasted with uranyl acetate, followed by Reynolds lead citrate, and imaged with the Hitachi HT7800 microscope (Hitachi, Japan) using the 20MP Xarosa camera (EMSIS, Germany).

#### Analysis of Transmission Electron Microscopy

Images were segmented to allow for the calculation of different mitochondrial parameters, such as size, aspect ratio, and number, using MitoNet, accessed through the empanada-napari interface^40^. Additionally, quadriceps TEM images were analyzed for the area of tubular aggregates using ImageJ2 (version: 2.16.0). Image analysis was performed blinded to genotype.

### Western blot analysis

Five to twenty micrograms of total tissue lysed in RIPA buffer (1% Triton X-100, 1% sodium deoxycholate, 0.1% SDS, 150 mM NaCl, 50 mM Tris HCl (pH 7.8), 1 mM EDTA, and 1 mM EGTA with Complete protease inhibitor cocktail (Roche)) on ice for 20min and centrifuged at 14,000rpm for 10min at 4°C. Alternatively, crude mitochondria were resuspended in 4x commercial Laemmli buffer and run in Bolt^TM^ 4-12% bis-tris Plus protein gels (Thermo Fisher, MA, USA, Cat. no. NW00120BOX) using MES running buffer. Proteins were transferred to nitrocellulose membranes using the iBlot™ 3 Transfer Stacks system (Thermo Fisher, MA, USA, Cat. No. IB31001). Standard detection methods for immunodetection using chemiluminescence Immun-Star HRP Luminol/Enhancer (Bio-Rad) were performed using the ChemiDoc XRS+ system (Biorad) or Amersham ImageQuant 800 (Cytiva) Western blot imaging system. All primary antibodies used are listed in Table 2. Band intensities were quantified using ImageJ (version: 2.16.0)

### BN-PAGE, in-gel activity assays

Mitochondria (100-150µg) isolated from mouse tissues were lysed in solubilization buffer [20 mM Tris-HCl (pH 7.4), 0.1mM EDTA, 50 mM NaCl and 10% (v/v) glycerol] containing either 1% (w/v) DDM or Digitonin and mixed with loading dye (5% Coomassie Brilliant Blue G-250, 150 mM Bis-Tris and 500 mM 6-aminocaproic acid pH 7.0). BN-Page and subsequent in-gel activity assays for complex I and complex IV were carried out as previously described^32^.

### Proteomic analysis

Snap-frozen total tissue from the apex of the heart and anterior quadriceps were used for mass spectrometry analysis.

#### Protein Digestion

Hearts were crushed using mortar and pestle in liquid nitrogen into a fine powder. The tissue powder was dissolved in 4% SDS in 100mM HEPES pH=8.5 (Volume / tissue ratio = 0.5 µL/µg). Proteins were reduced (10 mM TCEP) and alkylated (20 mM CAA) in the dark for 45 min at 45 °C. 25 µg protein was subjected to an SP3-based digestion^73^. Washed SP3 beads (SP3 beads (Sera-Mag(TM) Magnetic Carboxylate Modified Particles (Hydrophobic, GE44152105050250), Sera-Mag(TM) Magnetic Carboxylate Modified Particles (Hydrophilic, GE24152105050250) from Sigma Aldrich) were mixed equally, and 3 µL of bead slurry were added to each sample. Acetonitrile was added to a final concentration of 50% and washed twice using 70 % ethanol (V=200 µL) on an in-house manufactured magnet. After an additional acetonitrile wash (V=200µL), 5 µL digestion solution (10 mM HEPES pH 8.5 containing 0.5µg Trypsin (Sigma) and 0.5µg LysC (Wako)) was added to each sample and incubated overnight at 37°C. Peptides were desalted on a magnet using 2 × 200 µL acetonitrile. Peptides were eluted in 10 µL 5% DMSO in LC-MS water (Sigma Aldrich) in an ultrasonic bath for 10 min. Formic acid and acetonitrile were added to a final concentration of 2.5% and 2%, respectively. Samples were stored at - 20°C before subjection to LC-MS/MS analysis.

#### Liquid Chromatography and Mass Spectrometry

LC-MS/MS instrumentation consisted of an Easy-LC 1200 (Thermo Fisher Scientific) coupled via a nano-electrospray ionization source to an Exploris 480 mass spectrometer (Thermo Fisher Scientific, Bremen, Germany). An Aurora Frontier column (60cm length, 1.7 µm particle diameter, 75 µm inner diameter, IonOpticks) using a binary buffer system (A: 0.1 % formic acid and B: 0.1 % formic acid in 80% acetonitrile) based gradient was utilized as follows at a flow rate of 185 nL/min; a linear increase of buffer B from 4% to 28% within 100 min, followed by a linear increase to 40% within 10 min. The buffer B content was further ramped to 50 % within 4 minutes and then to 65 % within 3 minutes. 95 % buffer B was kept for a further 3 min to wash the column. The RF Lens amplitude was set to 45%, the capillary temperature was 275°C and the polarity was set to positive. MS1 profile spectra were acquired using a resolution of 30,000 (at 200 m/z) at a mass range of 450-850 m/z and an AGC target of 1 × 10^6^.

For MS/MS independent spectra acquisition, 34 equally spaced windows were acquired at an isolation m/z range of 7 Th, and the isolation windows overlapped by 1 Th. The fixed first mass was 200 m/z. The isolation center range covered a mass range of 500-740 m/z. Fragmentation spectra were acquired at a resolution of 30,000 at 200 m/z using a maximal injection time setting of ‘auto’ and stepped normalized collision energies (NCE) of 24, 28, and 30. The default charge state was set to 3. The AGC target was set to 3e6 (900% - Exploris 480). MS2 spectra were acquired in centroid mode. FAIMS was enabled using an inner electrode temperature of 100°C and an outer electrode temperature of 90°C. The compensation voltage was set to - 45V.

#### Data Analysis

Raw files were analyzed using Spectronaut v. 20.3.251119.92449^74^ In direct DIA mode using the Uniprot Mus Musculus (UP000000589 one-fasta-per-gene, 21783 protein sequences). Trypsin/P was selected as the cleavage rule using a specific digest type. The minimal peptide length was set to 7 and a total of 2 missed cleavages were allowed. The peptide spectrum match (PSM), peptide, and protein group FDR were controlled to 0.01. The mass tolerances were used with default settings (Dynamic,1). The directDIA +(deep) workflow was selected, and cross-run normalization was enabled. The protein group file was exported and LFQ intensities (MaxLFQ algorithm^75^) were log_2_ transformed. Statistically significantly different proteins were identified using a two-sided t-test followed by a permutation-based FDR calculation (s0=0.1, number permutations=500, FDR< 0.05) using Instant Clue^76^. The principal component analysis was performed after quantile normalization. Proteins with FDR < 0.05 were considered significantly regulated. Overrepresentation KEGG analyses were performed using ClusterProfiler (enrichKEGG) in R (Version 2024.09.1+39) on significantly differentially expressed proteins (FDR < 0.05) using all detected proteins as background. For enrichment analyses comparing 3 and 6-month DKO samples, Gene set enrichment analyses (GSEA) were performed using ClusterProfiler (gseKEGG) in R (Version 2024.09.1+39) on ranked proteins,and pathway enrichment was quantified using normalized enrichment scores (NES)^77^.

### Targeted lipidomic analysis

Twelve to eighteen milligrams of snap-frozen left ventricle of the heart and anterior quadriceps from male 6-month-old mice were homogenized using 1 × 2 mm ZrO beads (Techtum, Cat. no. ZrOB20) and 10 mg 1mm ZrO beads (Techtum, Cat. no. ZrOB10) in ice-cold isopropanol using the Fisherbrand Bead Mill 24 tissue homogenizer. After 15 minutes of sonication on ice and centrifugation, 150 µL of supernatant was transferred to LC-MS vials and placed in the autosampler for injection into Acquity Premier CSH C18 1.7µm 2.1 × 100 mm columns (Waters, Cat. no. 186009464). Sphingolipids were measured using an in-house prepared internal standard mix using the Waters Xevo TQ-S Triple Quadrupole Mass Spectrometer (Waters, Cat. no. WAA769). Positive, negative and cardiolipins were measured using Deuterated Lipidomics MaxSpec Mixture (Cayman Chemicals, Cat. no 40974) and cardiolipin internal standard (Avanti Lipids) with the Waters Xevo TQ-Sµ Triple Quadrupole Mass Spectrometer (Waters, Cat. no. QEG1182). Targetlynx files were generated for peak integration and quantification. Lipid abundances were normalized to tissue weight. For final analysis, ratios were calculated as the area under the curve of the compound (AUC) normalized to the AUC of its corresponding internal standard in arbitrary units (A.U.). Lipid classes included in the analysis are shown in Table 3. In total, 390 lipid species were measured, and 232 were detected.

**Table 3:** Measured lipid classes.

| Group | Lipid Class | Abbreviations |
| --- | --- | --- |
| Sphingolipids | Sphingolipids | SM, Cer, HexCer |
| Glycerophospholipids | Phosphatidylethanolamine | PE |
|  | Phosphatidylcholine | PC |
|  | Phosphatidylinositol | PI |
|  | Phosphatidylglycerol | PG |
|  | Cardiolipin | CL |
| Glycerolipids | Triacylglycerol | TG |
|  | Diacylglycerol | DG |
| Lysoglycerophospholipids | Lysophosphatidylcholine | LPC |
|  | Lysophosphatidylethanolamine | LPE |
| Fatty Acyls | Fatty Acids | FA |

#### Lipidomics data analysis

Lipidomics datasets were analyzed using LipidSigR in RStudio (Version 2024.09.1+39)^78,79^. Data were pre-processed by removing features with more than 70% missing values. Remaining features were normalized to the total lipid signal and log10-transformed. Statistical analysis was performed using two-sided t-tests with Benjamini-Hochberg correction, and lipid species with a false discovery rate (FDR) < 0.05 were considered significantly altered. Principal component analysis (PCA) was performed for dimensionality reduction. Lipid set enrichment analysis and pathway activity scoring were performed using LipidSigR based on significantly altered lipid species (FDR < 0.05 and |log_2_ fold change| ≥ 0.5), using all detected lipid species as background.

### RNA extraction and RT-qPCR

Snap-frozen tissue was used to isolate total RNA using TRIzol (Thermo Fisher Scientific), RNA integrity and concentration were determined using a Nanodrop spectrophotometer. Next, cDNA was synthesized using the High-Capacity cDNA Reverse Transcription kit (Applied Biosystems, Life Technologies). The QuantStudio 6 (Thermo Fisher Scientific), utilising TaqMan probes (Table 4) and TaqMan Universal Master Mix II (Life Technologies) was used for RT-qPCR. *β-Actin* was used as a housekeeping gene for normalization.

**Table 4:** List of TaqMan probes.

| GENE | TagMan Probe ID |
| --- | --- |
| *Mthfd2* | Mm00485276_m1 |
| *B-Actin* | Mm01205647_g1 |
| *Nd1* | Mm04225274_s1 |
| *Nd4* | Mm04225294_s1 |
| *18s rRNA* | Mm03928990_g1 |
| *Atf4* | Mm00515325_g1 |
| *Atf5* | Mm04179654_m1 |
| *Chop or Ddit3* | Mm01135937_g1 |
| *Pax7* | Mm01354484_m1 |
| *Myod1* | Mm00440387_m1 |
| *Nppa* | Mm01255747_g1 |
| *Nppb* | Mm01255770_g1 |

### Primary Satellite Cell Experiments from Mouse Skeletal Muscle

#### Cell culture

Satellite cells were isolated as previously described^80^. Clean myofibers from extensor digitorum longus (EDL) muscle were plated in proliferation medium [68% (v/v) DMEM + Glutamax (Gibco, Waltham, MA, United States), 20% (v/v) FBS (Gibco, Waltham, MA, United States), 10% (v/v) horse serum (HS) (Gibco, Waltham, MA, United States), 1% (v/v) chicken embryo extract (CEE) (ThermoFisher, Waltham, MA, United States), 0.0001% (v/v) fibroblast growth factor (FGF) (Prepotech, Rocky Hill, NJ, United States), 1% (v/v) penicillin-streptomycin (PS) (Sigma-Aldrich, Spruce St, MO, United States)] in a Petri dish precoated with Matrigel (Sigma-Aldrich, Spruce St, MO, United States) [1:10 (v/v) in DMEM+Glutamax] for 72 hours to allow satellite cells activation and migration. Finally, myogenic cells were isolated and amplified in proliferation medium and plated in proliferation or differentiation medium [97.0% (v/v) DMEM + Glutamax (Gibco, Waltham, MA, United States), 1% PS (Sigma-Aldrich, Spruce St, MO, United States) and 2% HS] for successive experiments.

For down-regulation of CHCHD2 and CHCHD10, cells were transfected with Accell Mouse *Chchd2* (M-055524-01-0005) siRNA (Dharmacon) and Accell Mouse *Chchd10* (M-041806-00- 0005) siRNA (Dharmacon) for the mouse cell line (20 nM) for a 48-h incubation period. As negative controls, cells with siGENOME RISC-free siRNA (Dharmacon). The siRNAs were preincubated with Lipofectamine RNAiMAX (Thermo Fisher Scientific, 13778150) in Opti-MEM (1X) (Gibco 31985-062) for 10 min prior to transfection. Knockdown efficiency was confirmed by RT-qPCR.

For RT-qPCR, 1 µg of total RNA was converted into cDNA using the iScript Reverse Transcription Supermix (Bio-Rad). RT-qPCR was performed using the CFX384 Touch Real-Time PCR Detection System (Bio-Rad) and SYBR® Green Master Mix (Bio-Rad) using the primers listed in Table 5. *Gapdh* was amplified as an internal standard. Data were analyzed according to the 2−ΔΔCT method^81^.

**Table 5:**
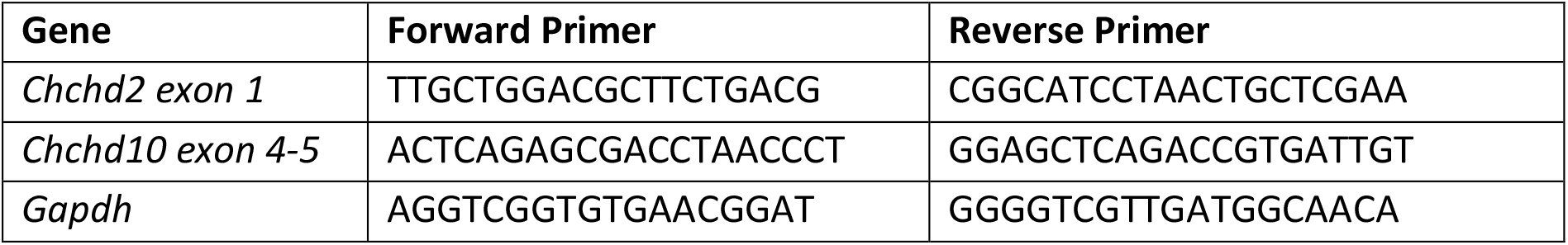
List of mouse qPCR primers.

#### Edu Cell Differentiation Assay

After transfection, primary cells were trypsinized, counted with Trypan Blue using the Countless II FL Automated Cell Counter (Invitrogen), and plated in Cell Carrier Ultra 96-well plates (PerkinElmer, 6055302). For the proliferation assay, cells were plated in proliferation medium at a density of 30,000 cells/cm². To assess proliferation, cells were pulsed with 10−6 M of EdU (ThermoFisher, C10640) in cell culture media for 2 h before fixation. Cells were fixed by adding 100 µL of 8% (w/v) paraformaldehyde to 100 µL of culture medium per well. Cells were subsequently washed three times with PBS for 5 min each. Then, 100 µL of 0.5% Triton X-100 in PBS was added to each well, and the plate was incubated at room temperature for 20 min. Cells were subsequently washed three times with PBS for 5 min each. Next, cells were incubated with the Click-iT reaction mixture for 30 min in the dark. Finally, cells were washed with PBS and incubated with Hoechst 33342 (H3570, Invitrogen) at 4 µg/mL with gentle shaking for 10 min. Images were acquired using the Operetta CLS High-Content Analysis System (PerkinElmer), with 5× Air/0.6 NA. Alexa Fluor 488 and Hoechst were excited with the 460-490 and 355-385 nm LEDs, respectively. The proliferation index was calculated as the ratio of EdU-positive to Hoechst-positive nuclei.

#### Myogenic differentiation assay

To induce myogenic differentiation and fusion, myoblasts were plated at high density (100,000 cells/cm2) onto Matrigel-coated Cell Carrier Ultra 96-well plates in the growth medium. Once adherent, cells were changed to differentiation medium for 24 and 48 hours. Cells were fixed by adding 100 µL of 8% (w/v) paraformaldehyde to 100 µL of culture medium per well, and the plate was transferred to an orbital shaker for 8 min. Cells were then washed 3 times with PBS for 5 min and permeabilized with 0.1% (v/v) Triton X-100/PBS shaking for 15 min. Cells were incubated with blocking solution (5% [v/v] FBS/PBS) shaking for 1 h. Myosin Heavy Chain (MyHC) antibody diluted in blocking solution (1:100) was incubated OV shaking at 4°C. Cells were washed 3 times with PBS for 5 min and the secondary antibody (Anti-Mouse IgG (H+L) Alexa 647; [1:1000]) was incubated shaking for 2 h in darkness. Cells were washed with PBS and incubated with Hoechst (H3570 Invitrogen) (4 μg/ml), shaking for 10 min. All steps were performed at RT. Images were acquired using the Opera Phenix CLS High-Content Analysis System (PerkinElmer), with 10× Air/0.6 NA. Alexa Fluor 647 and Hoechst were excited with the 620-650 and 355-385 nm LEDs, respectively. Automated image analysis was performed using Harmony Analysis Software (PerkinElmer). Image analysis is described in detail in the supplementary material.

#### Mitochondrial Membrane Potential Measurements

For mitochondrial membrane potential quantification, myogenic cells were incubated with Tetramethylrhodamine Ethyl Ester Perchlorate (TMRE; 50 nM) and MitoTracker Deep Red (MTDR; 100 nM). Nuclei were stained with NucBlue Live ReadyProbes Reagent (Thermo Fisher Scientific) at 1 drop per 10 mL of medium for 20 min at 37 °C and 5% CO₂. TMRE and MTDR signal intensities per cell were quantified using Harmony Analysis Software v4.9 (PerkinElmer) as previously described^82^. Image acquisition was performed with the Operetta CLS High-Content Analysis System (PerkinElmer) using a 63× water-immersion objective and excitation with the appropriate LED wavelengths. TMRE signal was normalized to mitochondrial mass (MTDR).

#### Mitochondrial Respiration of satellite cells

Mitochondrial respiration of satellite cells was analyzed using the Agilent Seahorse XFe96 Analyzer (Seahorse Biosciences), a microplate-based oxygen consumption system. Prior to the assay, the XFe96 microplate was precoated with Matrigel. The oxygen consumption rate (OCR) was measured in Agilent Seahorse XF Base Medium supplemented with 10 mM D-glucose, 1 mM sodium pyruvate, and 2 mM L-glutamine, with the pH adjusted to 7.4. Substrates and inhibitors were sequentially injected into the cartridge as follows: oligomycin (1 μM final), carbonyl cyanide m-chlorophenyl hydrazone (CCCP; 2 μM final), antimycin A (1 μM final), and rotenone (1 μM final). Respiratory parameters for OCR assays were calculated as previously described^83^. After the measurements were obtained, cell confluence was evaluated using the Incucyte® SX5 Live-Cell Analysis System (Sartorius) at 4× magnification to assess cell coverage, and confluence values were used for normalization.

### Statistical Analysis

Statistical analyses and graphs were plotted using GraphPad Prism v10 software (version 10.6.1) or R (R-Studio). All data are presented as means ± SD. Biological replicates (n) are denoted in the figure legends. For comparisons between two groups, two-sided unpaired Student’s t-tests were used unless otherwise stated. For multiple group comparisons, analysis of variance (ANOVA) with appropriate post hoc correction was applied. Values of p < 0.05 were considered statistically significant.

## AI Declaration

The authors used an AI-based language model to assist with improving the clarity of the manuscript text and for troubleshooting of code development. All scientific content, data analysis, and conclusions were generated and verified by the authors.

## Acknowledgments

We thank the Core Facility for Monoclonal Antibodies at the Helmholtz Institute, Munich (R. Feederle), for generating the custom CHCHD2 antibody. We acknowledge the Electron Microscopy Core Facility (L. Haag), the animal behavior core facility (Q.Yu), the Metabolic Phenotyping Centre (D. Rizo-Roca), and the Small-Molecule Mass Spectrometry Facility (A. Checa) at Karolinska Institutet for their support. We are grateful to E. Andersson (Karolinska Institutet) for providing access to the echocardiography system. We also thank D. Diehl (Max Planck Institute for Biology of Ageing) and Márcio Augusto Campos Ribeiro for expert assistance.

This study was supported by grants to R.F. from Vetenskapsrådet (2022-01477), Hjärnfonden, Parkinsonfonden, Åhlén-stiftelsen, StratNeuro, Åke Wiberg, Magnus Bergvalls Stiftelse, and Karolinska Institutet KID Funding Program for doctoral students.

## Author Contributions

Methodology: J.G., JD.HC., H.N., FC.N., JF.SR., D.A., D.M., and J.M. Analyses: J.G., JD.HC., H.N., FC.N., JF. SR., D.A., D.M., and R.F. Writing Original Draft: J.G. and R.F. Review & Editing: all authors. Visualization: J.G. and R.F. Supervision & Funding Acquisition: A.F., T.L., T.W., and R.F. Conceptualization: R.F.

## Data and materials availability

All data necessary to evaluate the conclusions in this paper are presented in the paper and/or in the supplementary materials.

---

**Fig. S1:**
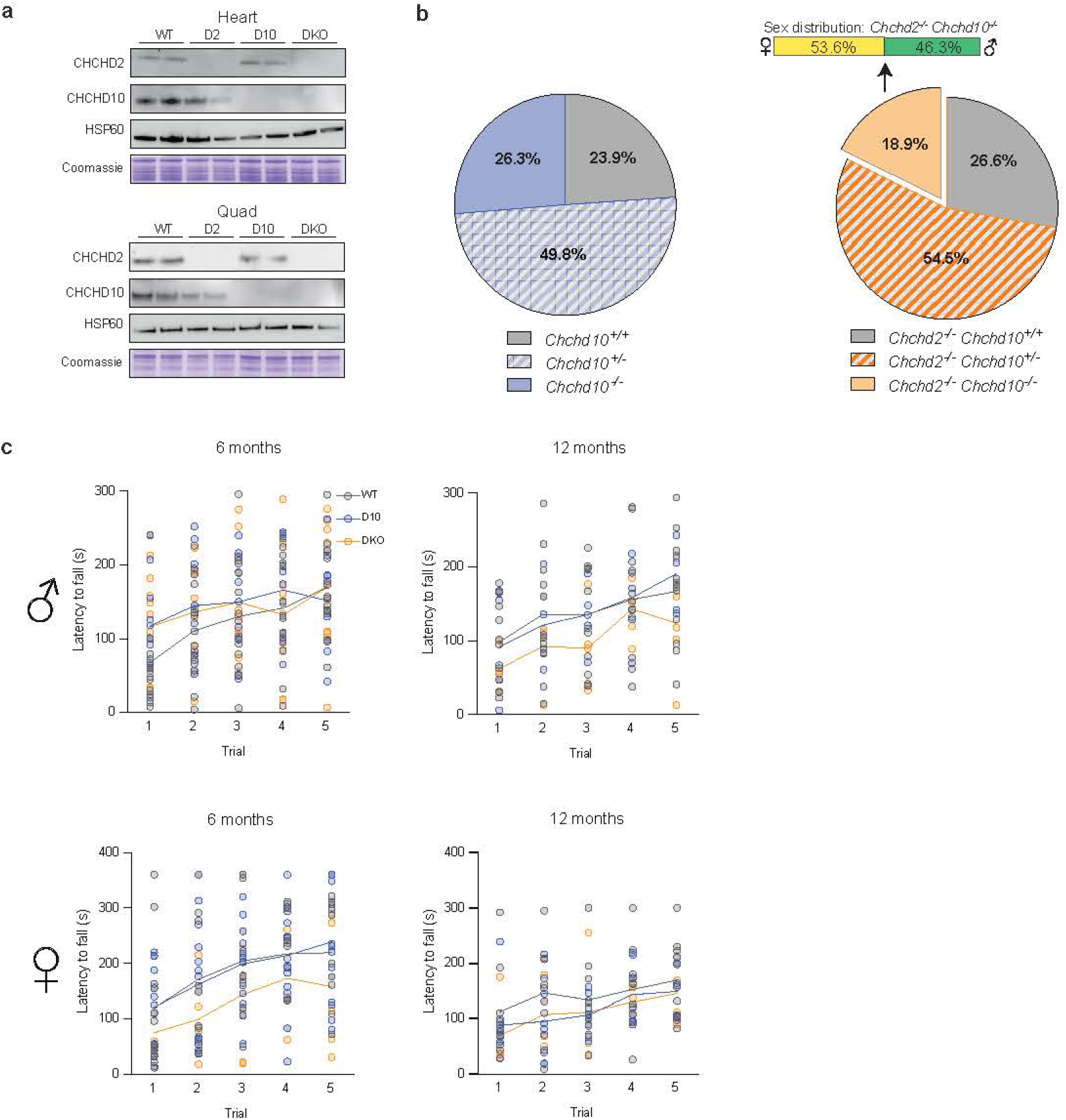
Generation and characterization of D10 KO and DKO mouse lines. **a,** Western blot showing CHCHD2 and CHCHD10 steady-state levels in mitochondria isolated from heart and quadriceps of WT, D2 KO, D10 KO, and DKO 3-month-old mice. HSP60 and Coomassie staining were used as loading controls. **b,** Mendelian distribution of offspring from heterozygous D10 KO crosses. For D10 KO mice: n=210 *Chchd10^+/-^*, n= 111 *Chchd10^-/-^,* n=101 *Chchd10^+/+^*. χ²(2)=0.483, p>0.05. Offspring from D2 KO and DKO mice were generated by crossing mice heterozygous for *Chchd10* and homozygous *Chchd2* knockout animals: n=127 *Chchd10^+/-^; Chchd2^-/-^;* n=44 *Chchd10^-/-^; Chchd2^-/-^*; n=62 *Chchd10^+/+^; Chchd2^-/-^.* χ²(2)=4.67, p=0.059. **c** Motor learning and endurance assessed as latency to fall (s) over five trials on the rod in WT, D10 KO, and DKO male and female mice at 6 and 12 months (n> 6 per genotype per timepoint).

**Fig. S2:**
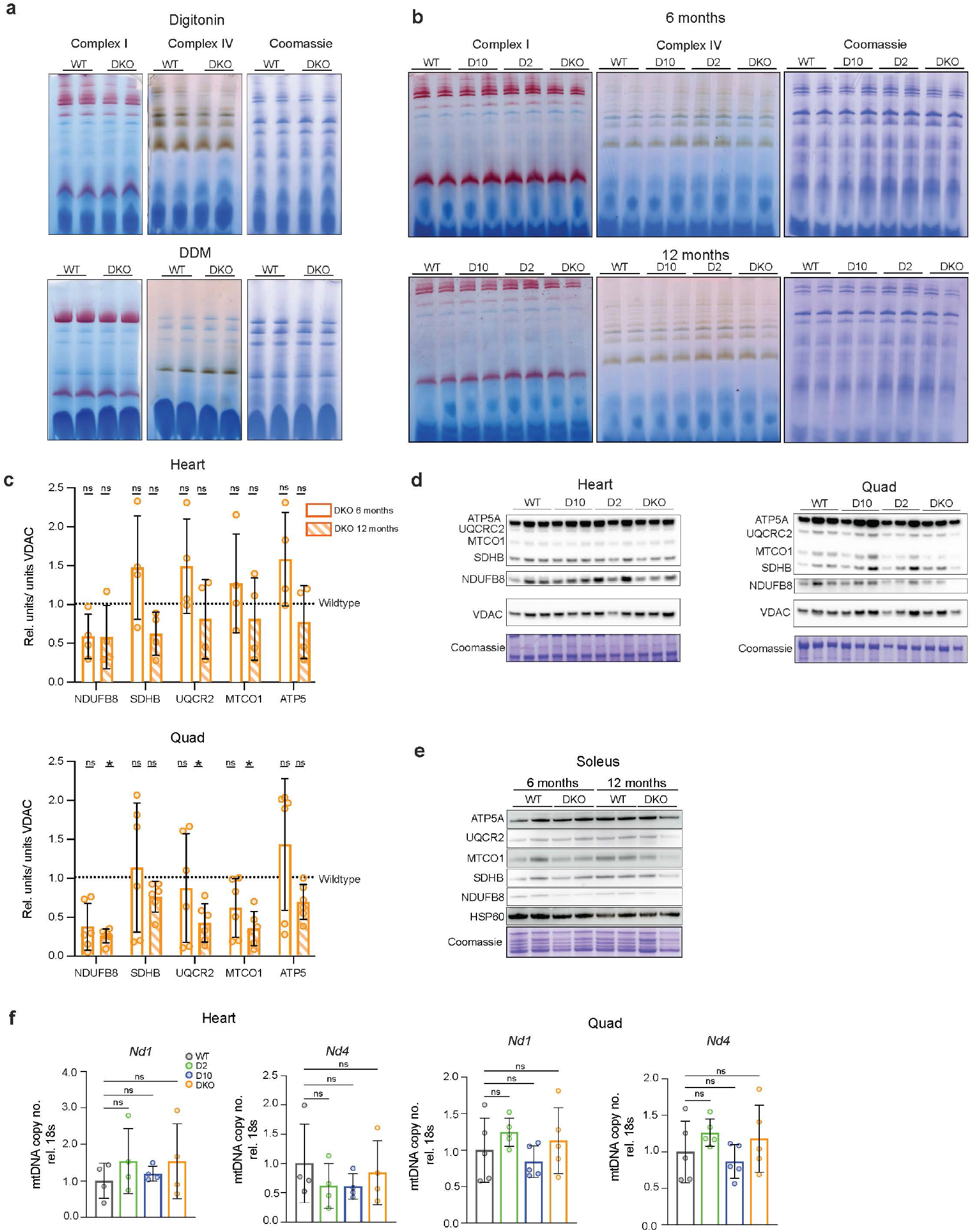
Impaired OXPHOS capacity in DKO mice is not driven by defects in individual complex assembly or by mtDNA depletion. Complex I and Complex IV in-gel activities in **a,** mitochondria isolated from quadriceps of 12-month-old mice using different detergent conditions (DDM and digitonin), and **b,** digitonin-solubilized mitochondria isolated from soleus from WT, D2 KO, D10 KO, and DKO at 6 and 12 months of age. **c,** Quantification of steady-state levels of OXPHOS subunits in mitochondria isolated from WT and DKO heart and quadriceps at 6 and 12 months of age, expressed relative to WT controls (dotted line). Data are presented as means ± SD (n> 4). *p <0.05; ns not significant. Western blot analyses of steady state OXPHOS subunits in **d,** the hearts and quadriceps of WT, D2 KO, D10 KO and DKO mice at 6 months, and **e,** soleus at 6 and 12 months of age. VDAC and HSP60 are used as loading controls, respectively. **f,** MtDNA copy number relative to 18S rRNA measured from hearts and quadriceps tissue at 6 months from WT, D2 KO, D10 KO and DKO mice. Data are presented as means ± SD; n> 4; ns not significant.

**Fig. S3:**
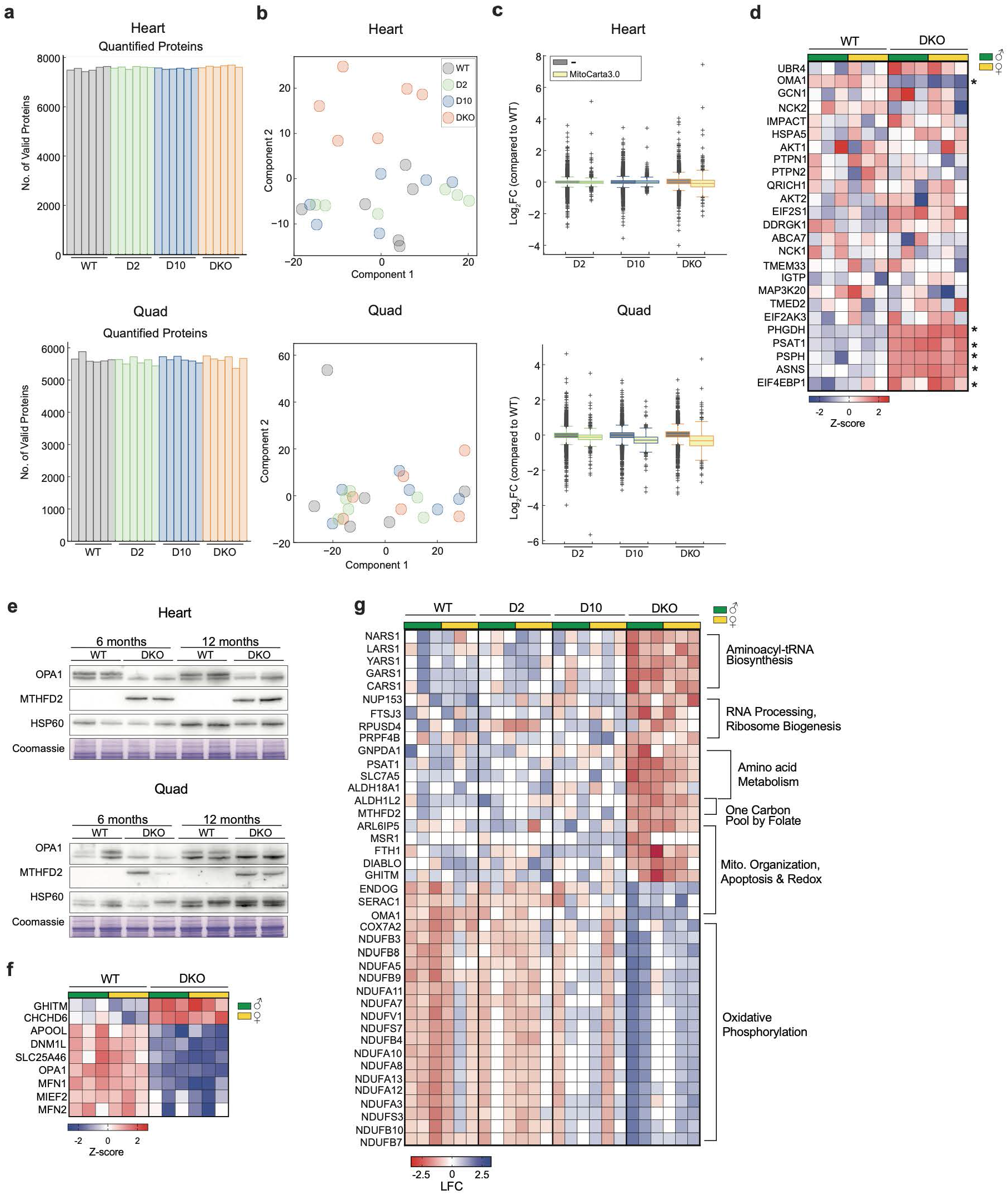
Proteomic profiling of heart and quadriceps lacking CHCHD2 and CHCHD10. **a,** Bar plots showing the total number of quan5ta5vely valid proteins iden5fied in each biological replicate from hearts and quadriceps of 6-month-old mice. **b,** Principal component analysis of heart and quadriceps proteomes. **c,** log_2_FC (DKO/WT) of mitochondrial proteins defined using the MitoCarta 3.0 database. **d,** Heatmap of heart proteomics showing proteins associated with the ISR. Sta5s5cally significant proteins are indicated by an asterisk. Each column represents a single mouse. **e,** Western blot analysis of MTHFD2 and OPA1 cleavage in total heart and quadriceps lysates from 6 and 12-month-old mice. HSP60 and Coomassie were used as loading controls. **f,** Heatmap of heart proteomics showing proteins involved in mitochondrial dynamics. **g,** Heatmap of top 50 DEPs quadriceps proteins analyzed by ANOVA analysis of WT, D2 KO, D10 KO and DKO.

**Fig. S4:**
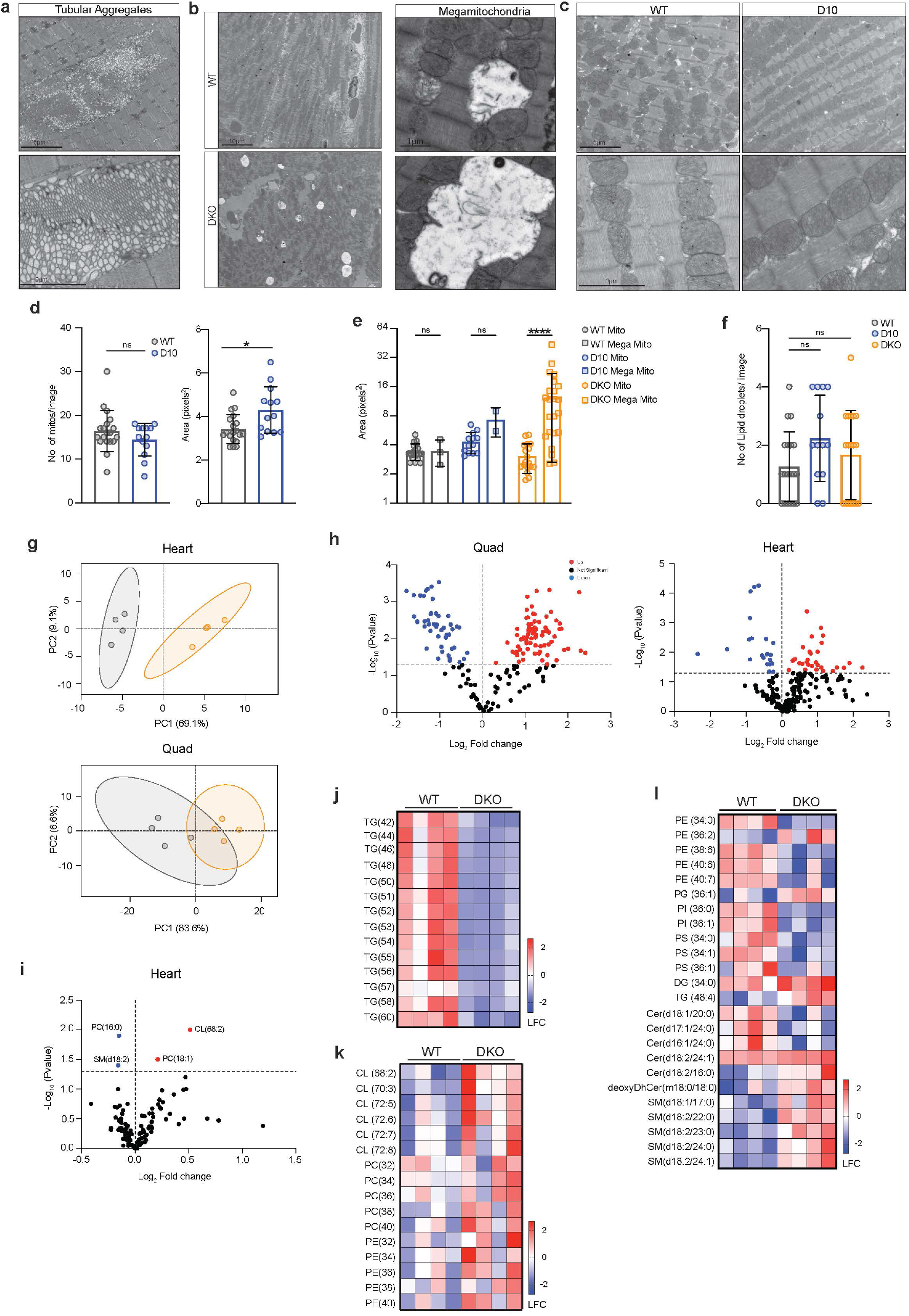
Alteration in mitochondrial morphology and lipid handling caused by CHCHD2-CHCHD10 deficiency. Representative TEM images from 6-month-old DKO quadriceps showing higher magnification tubular aggregates, and **b,** hearts showing low-magnification overview images and high-magnification images of megamitochondria. **c,** Representative TEM images from hearts of WT and D10 KO mice at 6 months of age. Scale bar: 5µm (low magnification) and 2µm (high magnification). **d,** Quantification of mitochondrial number and area from WT and D10 KO heart TEM images at 10,000X magnification. Data are presented as means ± SD; n= 3 mice, 4 images per animal. *p <0.05; ****p < 0.0001; ns not significant. Quantification of **e,** the area of healthy mitochondria versus megamitochondria, and **f,** lipid droplet count from heart TEM images of 6-month-old WT, D10 KO and DKO mice. Data presented as means ± SD; n> 2. ****p < 0.0001; ns not significant. **g,** PCA plot of heart and quadriceps lipidomics data from 6-month-old male mice. **h,** Volcano plots (log_₂_ fold change versus −log_₁₀_(FDR-adjusted P value)) of heart and quadriceps lipidomes from 6-month-old male mice. Significantly increased lipids are indicated in red, decreased lipids are in blue while unchanged lipids are denoted in black. **I,** Volcano plot (log_₂_ fold change versus −log_₁₀_(FDR-adjusted P value)) showing lipidomic changes from 12-month-old heart from D10 KO mice. **j-k,** Heatmap showing the log_2_ fold change of significantly changed lipid species from quadriceps of 6-month-old WT and DKO mice. Each column represents an individual animal. **I,** Heatmaps showing the log2 fold change of significantly changed lipid species from heart of 6-month-old WT and DKO mice. Each column represents an individual animal.

**Fig. S5:**
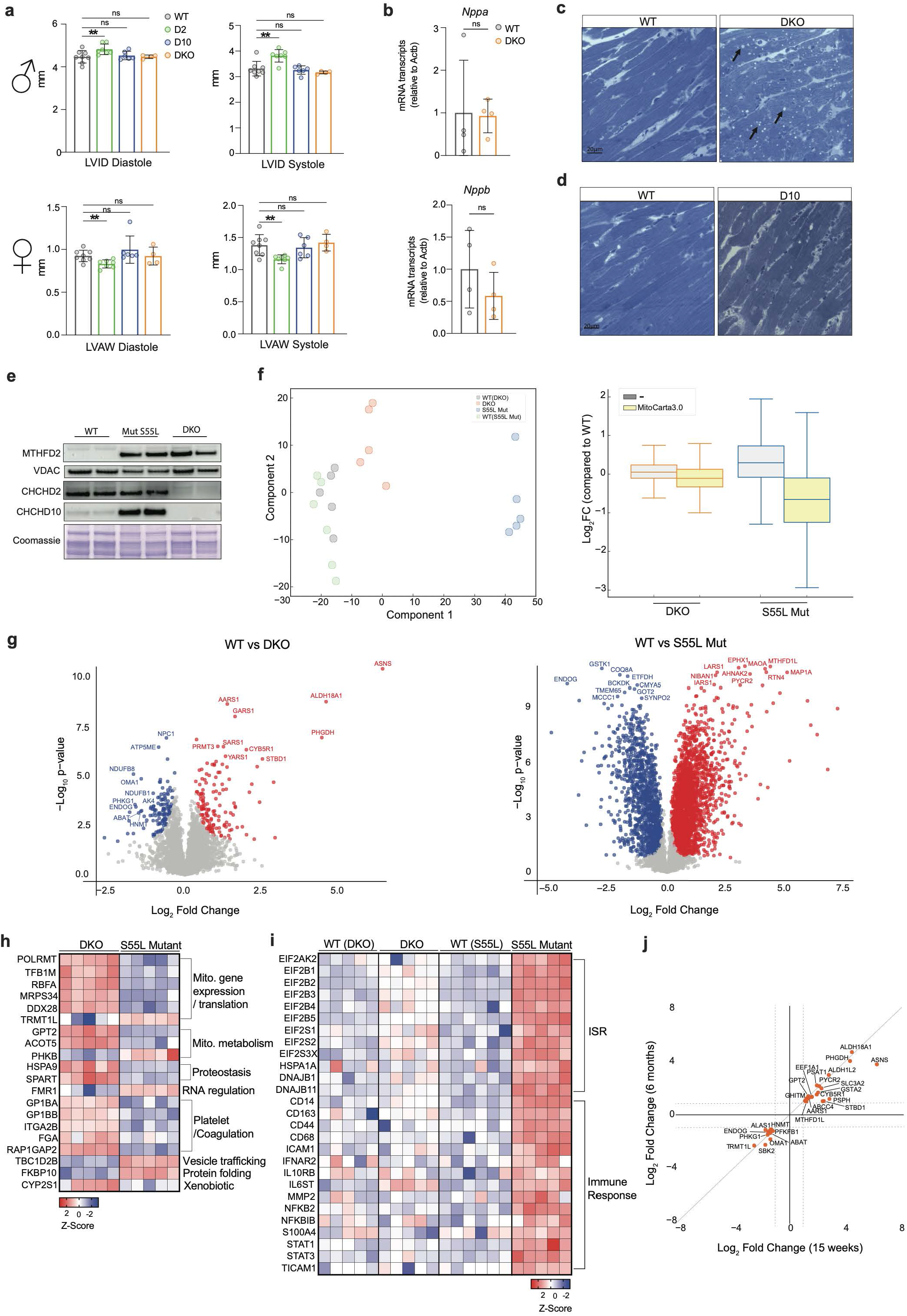
Comparative analysis of cardiac phenotypes in DKO, D2 KO, and CHCHD10 S55L mutant mice. **a,** Left ventricular internal diameter and left ventricular anterior wall quantification during systole and diastole of 6-9-month-old male and female WT, D2 KO, D10 KO and DKO mice. Data presented as means ± SD; n > 4. **p < 0.01; ns not significant. **b,** Relative mRNA transcript level of *Nppa* and *Nppb* of left ventricle from WT, D2 KO, D10 KO and DKO mice at 6 months. Data presented as means ± SD; n > 3. ns not significant. Representative toluidine blue staining of **c,** WT and DKO heart at 6 months of age. Black arrows indicate abnormal structures. **d,** of WT and D10 KO hearts at 6 months of age. **e,** Western blot analysis of MTHFD2, CHCHD2, and CHCHD10 in mitochondria isolated from the heart of WT, CHCHD10 S55L mutant, and DKO mice. VDAC was used as a loading control. **f,** Principal component analysis and log_2_FC of mitochondrial proteins defined using the MitoCarta 3.0 database of heart proteomes from 3-month-old DKO and S55L mutant mice. **g,** Volcano plot (log₂ fold change versus −log₁₀(P value)) of proteomes from 3-month-old DKO and S55L mutant hearts. Significantly upregulated proteins are denoted in red, downregulated proteins in blue, and unchanged in gray. Each dot represents a single protein. Proteins with |log_2_ fold change| ≥ 1 and FDR <0.05 were considered significant. **h,** Heatmap of log_2_ fold change of significant (FDR < 0.05) inverse differentially expressed proteins between DKO and S55L mutant (compared to their relative WT) heart at 3 months of age. **I,** Heatmap of log_2_ fold change of selected integrated stress response and immune response proteins in DKO and S55L mutant hearts with their relative WT. **j,** Log₂ fold change of 6-month-old versus log₂ fold change of 3-month-old DKO hearts, showing the top 40 significant (|Log₂ fold change| ≥ 1, FDR <0.05) proteins common to both timepoints.

**Fig. S6:**
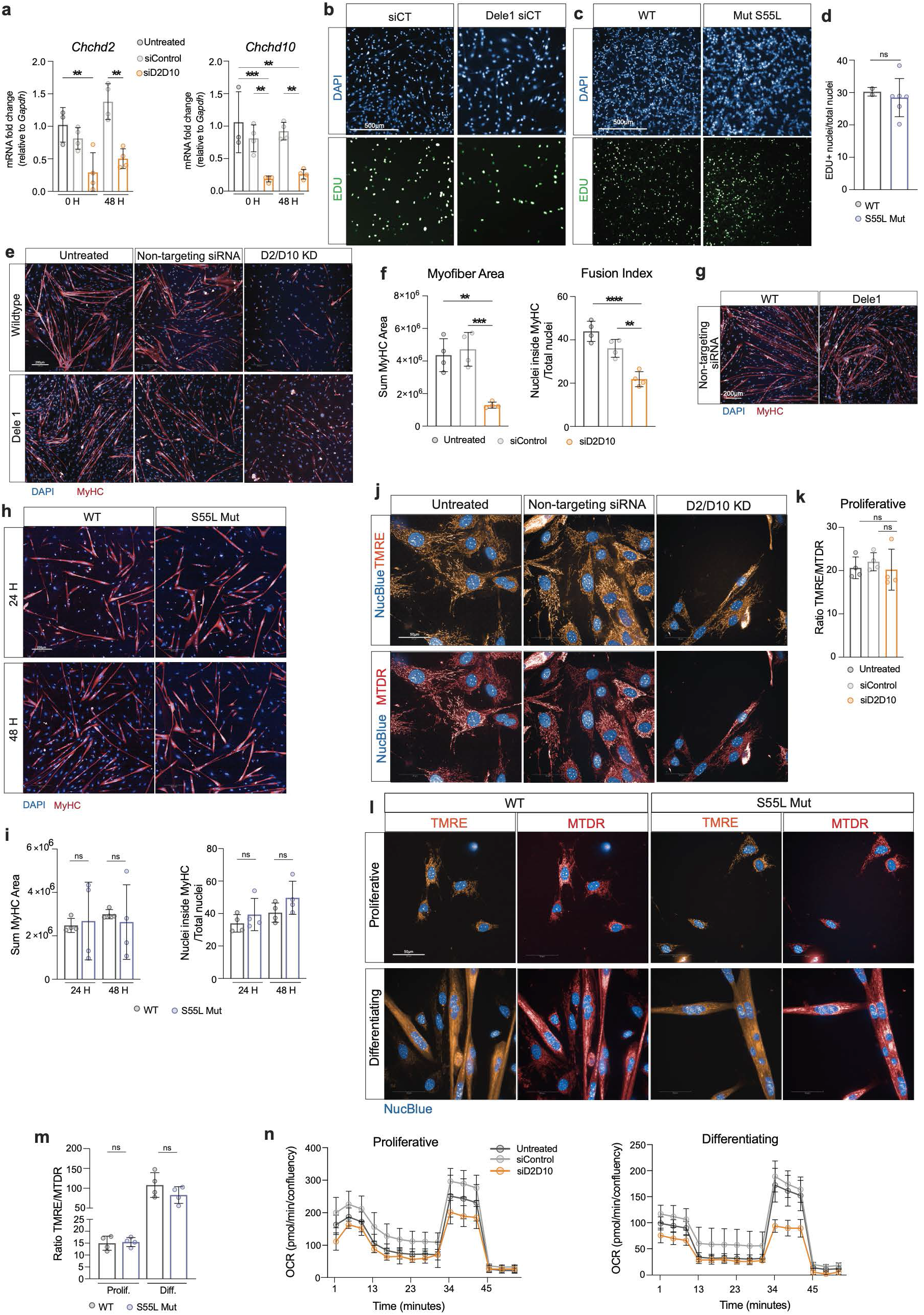
CHCHD2-CHCHD10 depletion impairs satellite cell proliferation, differentiation, and mitochondrial function. **a,** Relative mRNA transcripts of *Chchd2* and *Chchd10* in primary satellite cells isaolted form WT mice and treated with siControl and siD2/D10 at 0H and 48H after transfection, relative to *Gapdh*. Data are presented as means ± SD; n = 4; **p < 0.01; ***p < 0.001; **b,** Representative images of proliferative satellite cells isolated from WT and *Dele1* KO mice treated with siControl, stained for DAPI and Edu. Scale bar: 500µm. **c,** Representative images of proliferative satellite cells isolated from WT and S55L mutant and stained for DAPI and Edu. Scale bar: 500µm. **d,** Quantification of Edu+ nuclei of labeled primary satellite cells from WT and S55L mutant mice. Data are presented as means ± SD; n = 4; ns, not significant.. **e** Representative images of differentiating satellite cells isolated from WT mice and transfected with either siControl or siD2/D10, 24 h after induction of differentiation. Cells were stained with DAPI and MyHC. Scale bar:200µm. **f,** Quantification of myofiber area and fusion index after 24H of differentiation of WT, satellite cells treated with siD2/D10. Data presented as means ± SD; n = 4; **p < 0.01; ***p < 0.001. **g,** Representative images of WT and *Dele1* KO satellite cells 48 h after induction of differentiation and treatment with siControl. Cells were stained with DAPI and MyHC. Scale bar, 200 μm.**h,** Representative images of differentiating WT and CHCHD10 S55L mutant satellite cells at 24 h and 48 h after induction of differentiation. Cells were stained with DAPI and MyHC. Scale bar, 200 μm. **i,** Quantification of myofiber area and fusion index in WT and CHCHD10 S55L mutant satellite cells at 24 h and 48 h after induction of differentiation. Data are presented as mean ± SD; *n* = 4; ns, not significant. **j,** Representative images of proliferating WT satellite cells treated with siControl or siD2/D10 and stained with TMRE and MitoTracker Deep Red (MTDR). Scale bar, 50 μm. **k,** Quantification of the TMRE-to-MTDR fluorescence ratio in proliferating WT satellite cells treated with siControl or siD2/D10. Data are presented as mean ± SD; *n* = 4;. ns, not significant**. i,** Representative images of proliferating and differentiating WT and CHCHD10 S55L mutant satellite cells stained with TMRE and MitoTracker Deep Red (MTDR). Scale bar, 50 μm. **m,** Quantification of the TMRE-to-MTDR fluorescence ratio in proliferating and differentiating WT and CHCHD10 S55L mutant satellite cells. Data are presented as mean ± SD; *n* = 4; ns, not significant **n,** Oxygen consumption rate (OCR) measurements over time in proliferating and differentiating WT satellite cells treated with siControl or siD2/D10. *n* = 4.

